# Predicting the Effects of Per- and Polyfluoroalkyl Substance Mixtures on Peroxisome Proliferator-Activated Receptor Alpha Activity *in Vitro*

**DOI:** 10.1101/2021.09.30.462638

**Authors:** Greylin Nielsen, Wendy J. Heiger-Bernays, Jennifer J. Schlezinger, Thomas F. Webster

## Abstract

Human exposure to per- and polyfluoroalkyl substances (PFAS) is ubiquitous, with mixtures of PFAS detected in drinking water, food, household dust, and other exposure sources. Animal toxicity studies and human epidemiology indicate that PFAS may act through shared mechanisms including activation of peroxisome proliferator activated receptor α (PPARα). However, the effect of PFAS mixtures on human relevant molecular initiating events remains an important data gap in the PFAS literature. Here, we tested the ability of modeling approaches to predict the effect of diverse PPARα ligands on receptor activity using Cos7 cells transiently transfected with a full length human PPARα (hPPARα) expression construct and a peroxisome proliferator response element-driven luciferase reporter. Cells were treated for 24 hours with two full hPPARα agonists (pemafibrate and GW7647), a full and a partial hPPARα agonist (pemafibrate and mono(2-ethylhexyl) phthalate), or a full hPPARα agonist and a competitive antagonist (pemafibrate and GW6471). Receptor activity was modeled with three additive approaches: effect summation, relative potency factors (RPF), and generalized concentration addition (GCA). While RPF and GCA accurately predicted activity for mixtures of full hPPARα agonists, only GCA predicted activity for full and partial hPPARα agonists and a full agonist and antagonist. We then generated concentration response curves for seven PFAS, which were well-fit with three-parameter Hill functions. The four perfluorinated carboxylic acids (PFCA) tended to act as full hPPARα agonists while the three perfluorinated sulfonic acids (PFSA) tended to act as partial agonists that varied in efficacy between 28-67% of the full agonist, positive control level. GCA and RPF performed equally well at predicting the effects of mixtures with three PFCAs, but only GCA predicted experimental activity with mixtures of PFSAs and a mixture of PFCAs and PFSAs at ratios found in the general population. We conclude that of the three approaches, GCA most accurately models the effect of PFAS mixtures on hPPARα activity *in vitro*.

**Highlights:**

- Perfluorinated carboxylic acids are full human PPARα agonists
- Perfluorinated sulfonic acids are partial human PPARα agonists
- GCA predicts human PPARα activity for mixtures of full and partial agonists
- GCA predicts human PPARα activity for mixtures of agonists and competitive antagonists
- GCA accurately predicts human PPARα activity in response to PFAS mixtures

## 1.0 Introduction

People are exposed to mixtures of per- and polyfluoroalkyl substances (PFAS) through water, air, food, household dust, soil, breast milk, and consumer products (Awad et al. 2020; Begley et al. 2008; De Silva et al. 2021; Death et al. 2021). Recent analyses showed the detection of 10 PFAS in more than 50% of treated drinking water samples in a targeted analysis in the United States, and the detection of 14 PFAS in more than 90% of dust samples at eight U.S. childcare centers (Boone et al. 2019; Zheng et al. 2020). Exposure to PFAS mixtures is documented in human biological samples with multiple PFAS ubiquitously detected in the U.S. general population over multiple cycles of the National Health and Nutrition Examination Survey (CDC 2019; Kato et al. 2011). Assessing health risks associated with exposure to PFAS mixtures remains a critical gap with implications for regulatory standards setting for drinking water and other exposure media.

Individual PFAS are associated with adverse health effects in human epidemiological studies supported by toxicity evaluations in animal models. Perfluorooctanoic acid (PFOA) and perfluorooctane sulfonic acid (PFOS) have received more research attention than other PFAS congeners, though studies with other PFAS, especially perfluorohexane sulfonic acid (PFHxS) and perfluorononanoic acid (PFNA), are rapidly increasing in number. Exposures to PFOS, PFOA, and PFHxS in sensitive human subpopulations are associated with suppression of the immune response (DeWitt et al. 2019; EFSA 2020; Pachkowski et al. 2019), which is supported by animal toxicity data (DeWitt et al. 2019). PFAS also exhibit metabolic toxicity. Cross-sectional and longitudinal studies show an association between multiple PFAS and elevated total and low density lipoprotein cholesterol levels in adults and children (Fenton et al. 2021; Nelson et al. 2010; Rappazzo et al. 2017). PFAS affect hepatic and whole body lipid metabolism in animal toxicity studies as well (Das et al. 2017; Pfohl et al. 2020; Rebholz et al. 2016; Zhang et al. 2018).

Mechanisms underlying the biological effects induced by PFAS are not completely understood. However, many PFAS activate several nuclear receptors involved in metabolic signaling including peroxisome proliferator-activated receptor alpha (PPARα), implicating nuclear receptor signaling as an important molecular initiating event (Behr et al. 2020a; Bijland et al. 2011; Bjork et al. 2011; Li et al. 2019; Rosen et al. 2017a; Schlezinger et al. 2020b). PPARα is a ligand activated transcription factor that plays a critical role in hepatic nutrient metabolism in humans by responding to endogenous and dietary lipids (Ferré 2004; Keller et al. 1993; Kersten 2014; Mandard et al. 2004). Different PFAS congeners activate PPARα in reporter assays with human and mouse PPARα (Behr et al. 2020a; Buhrke et al. 2013; Carr et al. 2013; Maloney and Waxman 1999; Rosenmai et al. 2018; Rosenmai et al. 2016; Shipley et al. 2004; Takacs and Abbott 2007; Vanden Heuvel et al. 2006; Wolf et al. 2014; Wolf et al. 2012; Wolf et al. 2008) (Table S1). PFAS also activate PPARα in human-like and human primary hepatocytes (Behr et al. 2020b; Bjork et al. 2011; Buhrke et al. 2013; Buhrke et al. 2015; Louisse et al. 2020; Rowan-Carroll et al. 2021), and in wild type (Marques et al. 2020; Rosen et al. 2008a; Rosen et al. 2008b) and humanized PPARα rodent models (Schlezinger et al. 2020b). In mouse models, PPARα regulates more than 75% of hepatic gene expression altered by PFOA, PFNA, PFOS, and PFHxS (Rosen et al. 2008a; Rosen et al. 2017b; Rosen et al. 2008b). Thus, PPARα activation is an important and human-relevant molecular initiating event shared across multiple PFAS congeners.

Few studies have examined the joint effect of PFAS mixtures under human-relevant conditions. Most use extreme endpoints such as cytotoxicity and organism lethality where the driving mechanism is likely dissimilar to mechanisms underlying health effects of PFAS at human relevant exposures (Ding et al. 2013; Flynn et al. 2019; Hoover et al. 2019; McCarthy et al. 2021; Ojo et al. 2020). Two studies using reporter assays with a mouse PPARα ligand binding domain-Gal4 system employed modeling approaches to compare predicted receptor activity to observed activity following PFAS mixture treatment (Carr et al. 2013; Wolf et al. 2014). Carr et al., 2013 employed a nonlinear additivity model to predict responses for fixed ratio mixtures and found that binary and complex PFAS mixtures had a less than additive effect on mouse PPARα activation. Wolf et al., 2014 employed both concentration additive and response additive approaches to predict the effect of PFAS mixtures on mouse PPARα activity. Their analysis used estimates of potency (a measure of receptor activity expressed in terms of the amount required to produce an effect of given intensity) and found that the effect of binary and multi-component PFAS mixtures could be predicted by both concentration and response addition at lower concentrations but departed from these additivity models at higher concentrations with the direction of the deviation related to the composition of the PFAS mixture.

There are important limitations to these previous approaches, however. First, both studies used a two hybrid (protein-DNA) reporter assay using only the ligand-binding domain of mouse PPARα in a Gal4 construct. This artificial system may reflect binding affinities of PFAS for PPARα but transcriptional efficacies may be misrepresented (Gearing et al. 1993; Smith and O’Malley 2004; Webster et al. 1988). Second, there are well characterized differences between mouse and human PPARα, and human PPARα is less responsive to PFAS than the mouse homolog (Gonzalez and Shah 2008; Takacs and Abbott 2007; Wolf et al. 2012; Wolf et al. 2008). Last, there are some previous indications that PFAS may activate PPARα with differing potency and efficacy (Behr et al. 2020a; Rosenmai et al. 2018; Rosenmai et al. 2016; Vanden Heuvel et al. 2006; Wolf et al. 2014). However, the relative potency factor model assumes mixture components elicit the same maximal receptor response (efficacy).

Therefore, in this study, we used full length human PPARα (hPPARα) and hypothesized that modeling approaches that account for differences in both PFAS potency and efficacy will predict hPPARα activity more accurately than other approaches. To test this hypothesis, we first compared three additivity models in Cos7 cells transiently transfected with full length hPPARα and a peroxisome proliferator response element-driven luciferase reporter. We generated concentration response curves for well-characterized full and partial hPPARα agonists and a competitive antagonist. We treated cells in a grid design with mixtures of hPPARα ligands and then predicted activity with three models: effect summation (ES), relative potency factor (RPF), and generalized concentration addition (GCA) with the latter using a concentration-response function based on a two-step pharmacodynamic model for a receptor with a single binding site. We then established concentration response curves for sixseven different PFAS and tested the ability of these modeling approaches to predict hPPARα activity for PFAS mixtures. Our analysis establishes GCA as an accurate model to predict hPPARα activation by mixtures of diverse ligands including PFAS.

## 2.0 Methods

### 2.1 Chemical Suppliers

The source, purity, and CAS registry number for hPPARα ligands (and their acronyms) are reported in Table 1. Stock solutions of GW7647, GW6471, pemafibrate, gemfibrozil, ETYA, and mono(2-ethylhexyl) phthalate (MEHP) were prepared in dimethyl sulfoxide (DMSO, CASRN 67-68-5, AmericanBio, Inc. (Canton, MA, USA). PFOA, PFNA, PFOS, PFHxS were dissolved in DMSO and diluted with NERL water (Thermo Fisher Scientific, Waltham, MA) to a stock solution in 50% DMSO:50% NERL water. NBP2 stock solutions were prepared in 50% DMSO:50% NERL water. Stock solutions of GenX were prepared in NERL water. All other reagents were from Thermo Fisher Scientific, unless noted.

**Table 1.** Chemical suppliers, purity, and numeric identifier for PPARα ligands.

| Chemical | CAS Registry Number | Purity | Source |
| --- | --- | --- | --- |
| 5,8,11,14-Eicosatetraynoic acid (ETYA) | 1191-85-1 | $\geq 97\%$ | Sigma-Aldrich |
| Gemfibrozil | 25812-30-0 | 98% | Toronto Research Chemicals Inc. |
| GW6471 | 880635-03-0 | $\geq 98\%$ | Sigma-Aldrich |
| GW7647 | 265129-71-3 | $\geq 96\%$ | Sigma-Aldrich |
| Mono-(2-ethylhexyl) phthalate (MEHP) | 4376-20-9 | 97% | Sigma-Aldrich |
| Pemafibrate (Pema) | 848259-27-8 | 98% | Toronto Research Chemicals Inc. |
| Perfluoroheptanoic acid (PFHpA) | 375-85-9 | $\geq 98\%$ | Santa Cruz Biotechnology |
| Perfluorooctanoic acid (PFOA) | 335-67-1 | 95% | Sigma-Aldrich |
| Perfluorononanoic acid (PFNA) | 375-95-1 | $\geq 97\%$ | Santa Cruz Biotechnology |
| Perfluorooctane sulfonic acid (PFOS) | 1763-21-1 | $\geq 97\%$ | Santa Cruz Biotechnology |
| Potassium-perfluorohexane sulfonate (PFHxS) | 3871-99-6 | $\geq 98\%$ | Santa Cruz Biotechnology |
| Nafion By-Product 2 (NBP2) | 749836-20-2 | 95% | Synquest Laboratories |
| Perfluoro(2-methyl-3-oxahexanoic) acid (GenX) | 13252-13-6 | $\geq 96\%$ | Santa Cruz Biotechnology |

### 2.2 Reporter Assays

Cos7 cells were maintained at 37°C with CO_2_ supplementation in Dulbecco’s Modified Eagle Medium (DMEM, Corning, Corning, NY) with 10% fetal bovine serum (FBS, Gemini Bio-Products, West Sacramento, CA) and antibiotic-antimycotic (100 U/mL penicillin, 100 μg/ml streptomycin, 0.25 μg/ml amphotericin B). On day one, cells were plated at 20,000 cells per well in 96-well, white-sided plates in 0.1 mL DMEM with 5% FBS. On day two, cells were transiently transfected with a full-length human *PPARA* expression vector (generously provided by Dr. Frank Gonzales (Sapone et al. 2000)) in addition to a peroxisome proliferator response element 3x-TK-Luc reporter construct (Plasmid 1015; Addgene; (Kim et al. 1998)), and a lab-made cytomegalovirus promoter-driven enhanced green fluorescent protein (GFP) reporter construct using Lipofectamine 2000 (Invitrogen, Carlsbad, CA, USA). On day three, medium was replaced with 0.2 mL unsupplemented DMEM, and cells received no treatment (medium only), vehicle (0.1% DMSO), the positive control (GW7647, 1 × 10^−6^ M), or hPPARα ligands alone or in combinations at varying concentrations (GW7647, 1 × 10^−12^ M to 1 × 10^−6^ M; Pemafibrate, 1 × 10^−13^ M to 1 × 10^−7^ M; MEHP 1 × 10^−8^ M to 4 × 10^−5^M; PFHpA, 1 × 10^−6^ M to 8 × 10^−5^ M; PFOA 1 × 10^−8^ M to 1 × 10^−4^ M; PFNA 1 × 10^−8^ M to 4 × 10^−5^ M; PFOS 4 × 10^−8^ M to 4 × 10^−5^ M; PFHxS 1 × 10^−7^ M to 1 × 10^−4^ M; NBP2 1 × 10^−8^ M to 1 × 10^−4^ M.) Concentrations were selected from range-finding experiments that identified the highest non-toxic concentration for each chemical. Toxicity was assessed by comparing the average GFP fluorescence in wells treated with positive control to wells treated with experimental chemicals; treatments that reduced fluorescence were eliminated from the analysis (data not shown). In each experimental plate, six wells were treated with the positive control and six wells were left untreated. At least two wells per plate were treated with vehicle or hPPARα ligands and mixtures. After 24 hours of treatment, cells were lysed using Steady-Luc Firefly HTS Reagent (Biotium, Freemont, CA, USA). Luminescence and fluorescence were measured on a Synergy2 plate reader (Biotek Inc, Winooski, VT, USA) following a five-minute incubation on a platform rocker. Four biological replicates were performed for each experiment.

Luminescence from each well was normalized by dividing by the GFP-fluorescence. The average normalized background signal in wells treated with medium only was subtracted from all experimental wells. The percent of maximum hPPARα activity was determined by dividing the background corrected signal from each well by the background positive control signal, which was identified as a maximally efficacious concentration of GW7647. The percent of maximum hPPARα activity was averaged across duplicate or triplicate wells. This normalization method is designed to minimize intra- and inter-experimental variation (Rajapakse et al. 2004).

### 2.3 Mixtures Experimental Design

We examined the predictive capacity of additivity models by performing experiments with binary mixtures of full hPPARα agonists (pemafibrate and GW7647), a full agonist and a partial agonist (pemafibrate and MEHP), and a full agonist and competitive antagonist (pemafibrate and GW6471). Cells were treated with increasing concentrations of each chemical in a grid design. We also performed experiments with binary mixtures of PFOA with PFOS and GenX with NBP2 in a grid design to confirm the applicability of predictive models to PFAS mixtures.

We then conducted a series of experiments with multicomponent PFAS mixtures in a ray design. We created an equipotent mixture of PFHpA, PFOA, and PFNA in which each chemical contributed equally to the total mixture potency. The total concentration of PFAS in the mixture was 1×10^−4^M in this stock solution of perfluorinated carboxylic acids (PFCAs). In this PFCA mixture, PFOA served as the reference chemical to determine the relative potency of the components. The PFCA stock solution was serially diluted to produce a range of test concentrations between 1×10^−7^ M to 1×10^−4^ M total PFAS. We created a similar stock solution with PFOS, PFHxS, and NBP2 in which each chemical contributed equally to mixture potency with PFOS as the reference chemical. This perfluorinated sulfonic acid (PFSA) stock mixture solution was serially diluted to provide test concentrations between 1×10^−7^M to 5×10^−5^M for total PFAS. Finally, we tested a mixture of PFOA, PFOS, PFNA, and PFHxS at the ratio of median concentrations found in the U.S. general population in the 2015-2016 NHANES cycle (CDC 2019). The ratio of chemicals in the mixture was 24:6:7.5:3 for PFOS:PFHxS:PFOA:PFNA. The highest concentration of total PFAS in this NHANES mixture was 1.2×10^−4^ M, and the stock solution was serially diluted to test concentrations between 1.2 × 10^−8^ M to 4.7×10^−5^ M total PFAS. For all experiments conducted in a ray design, two to three wells per experimental plate were treated with each tested concentration. Each experiment was performed at least four times.

### 2.4 Predictive Modeling and Statistical Analysis

Concentration response curves for each test chemical were plotted in GraphPad Prism (version 9.0.0) and goodness of fit was compared between three and four parameter Hill functions at the α=0.05 level of statistical significance. Individual curve fits were used to model the effects of chemical mixtures on hPPARα activation with three different predictive models including effect summation (ES), relative potency factor (RPF), and generalized concentration addition (GCA). ES models the combined response to mixtures as the sum of the effect for each compound at a given concentration. For a binary mixture, the predicted effect is given by

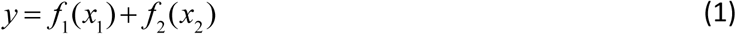

where *f*_i_(*x*_i_) is the concentration response for compound *i*. ES is expected to over-predict the empirical mixture response at high concentrations and is considered of limited utility in predicting the toxicity of chemical mixtures (Berenbaum 1989).

The relative potency factor approach is a special case of concentration addition where the concentration response curves for individual mixture components are assumed to differ only in potency. This approach assumes individual concentration response curves have the same shape and maximal effect and differ only in potency. RPF compares the potency of each chemical in the mixture to a reference compound; this approach is employed in the toxic equivalency factors used to assess additive effects of dioxin-like compounds (Van den Berg et al. 2006) and has been proposed as a method to model additive effects of PFAS on liver endpoints (Bil et al. 2021). The response predicted by RPF for a binary mixture is given by:

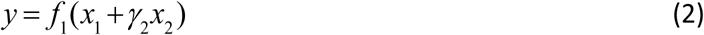

In Equation 2, f_1_ is the concentration response function for reference compound x_1_, and *γ*_2_ is the relative potency factor for the second compound with a concentration of x_2_. Typically, the relative potency factor is the EC_50_ for the reference compound divided by the EC_50_ for the second compound. RPF considers only differences in potency for mixture components and is expected to inaccurately predict effects when compounds differ in efficacy.

Some receptor agonists do not obey the stringent parallel concentration response curve requirement of RPF. We therefore developed generalized concentration addition which models the effect of chemical mixtures as a function of the potency and efficacy of each chemical in the mixture and can predict effects with mixtures of full and partial receptor agonists as well as agonists and competitive antagonists (Howard et al. 2010; Howard and Webster 2009; Schlezinger et al. 2020a; Watt et al. 2016). The general definition of GCA for a binary mixture (extendable to more compounds) is given by

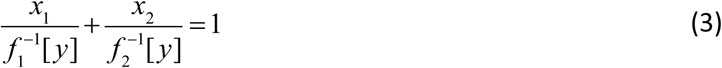

where the denominators are the inverse concentration-response functions of the compounds evaluated at effect level *y*, yielding real numbers (Howard and Webster 2009). The CA and RPF models are special cases of GCA. Rather than using arbitrary concentration-response functions, we use pharmacodynamic models (PDM). For a receptor like PPARα that has a single ligand binding site where activation can occur following binding, a 3-parameter Hill function with Hill coefficient (slope parameter) equal to 1 is appropriate.

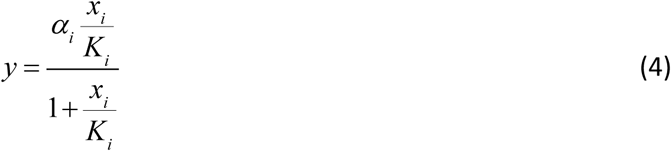

where α_i_ is the efficacy of each compound, *x*_i_ is the concentration of each compound, *K*_i_ is the EC_50_ for each compound and *y* is the predicted response. With GCA, the effect of a binary mixture is then predicted with Equation 5 (Howard and Webster 2009):

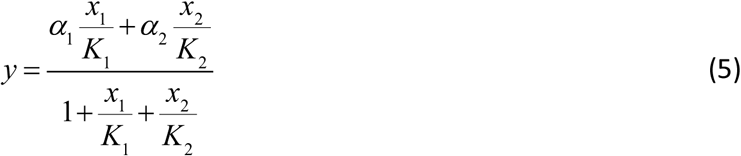

Application of GCA here yields the same result (equation 5) as a pharmacodynamic model of a mixture of receptor ligands. The Gaddum equation for a competitive antagonist is a special case of Equation (5) where the efficacy of the antagonist is zero (Howard and Webster 2009).

For simplicity in comparing ES, RPF and GCA, we used the three parameter Hill model for each compound (equation 4). No statistically significant difference was seen between three and four parameter Hill functions for any of the ligands tested in this analysis (Table S2). We used PFOA as the reference compound for the RPF model for PFCA mixtures and mixtures of PFCAs and PFSAs. PFOS was the reference compound for mixtures of PFSAs.

For grid design mixtures experiments, predicted response surfaces were plotted in R version 4.0.2 (2020-06-22). Experimental data from ray design experiments were plotted in GraphPad Prism (version 9.0.0) and were compared with responses predicted by the three modeling approaches. Concordance between experimental mixtures data and predicted response were compared statistically by calculating root mean square error (RMSE). Lower RMSE values indicate a better fit between predicted and empirical data.

## 3.0 Results

### 3.1 hPPARα Reporter System Functions Appropriately with Known Full and Partial Agonist

We first tested the hPPARα reporter system by establishing concentration response curves for known full and partial hPPARα agonists. Concentration-response curves for two full hPPARα agonists (GW7647 and pemafibrate) and three partial hPPARα agonists (MEHP, gemfibrozil, ETYA) are shown in Figure 1. The concentration-response curves for GW7647, pemafibrate, and MEHP were well-fit with a three-parameter Hill function with Hill slope of 1; curve fits were not significantly different from fits achieved with a four-parameter Hill function where the slope of the curve was not constrained to 1 (Table S2). Both full agonists induced maximal hPPARα activity and had similar potencies (an EC_50_ of 1.8×10^−11^ M for GW7647 compared to an EC_50_ of 2.3×10^−11^ M for pemafibrate). MEHP was less potent and efficacious than the full agonists with a maximum efficacy of 60% and EC_50_ of 5.2×10^−6^ M.

**Figure 1.**
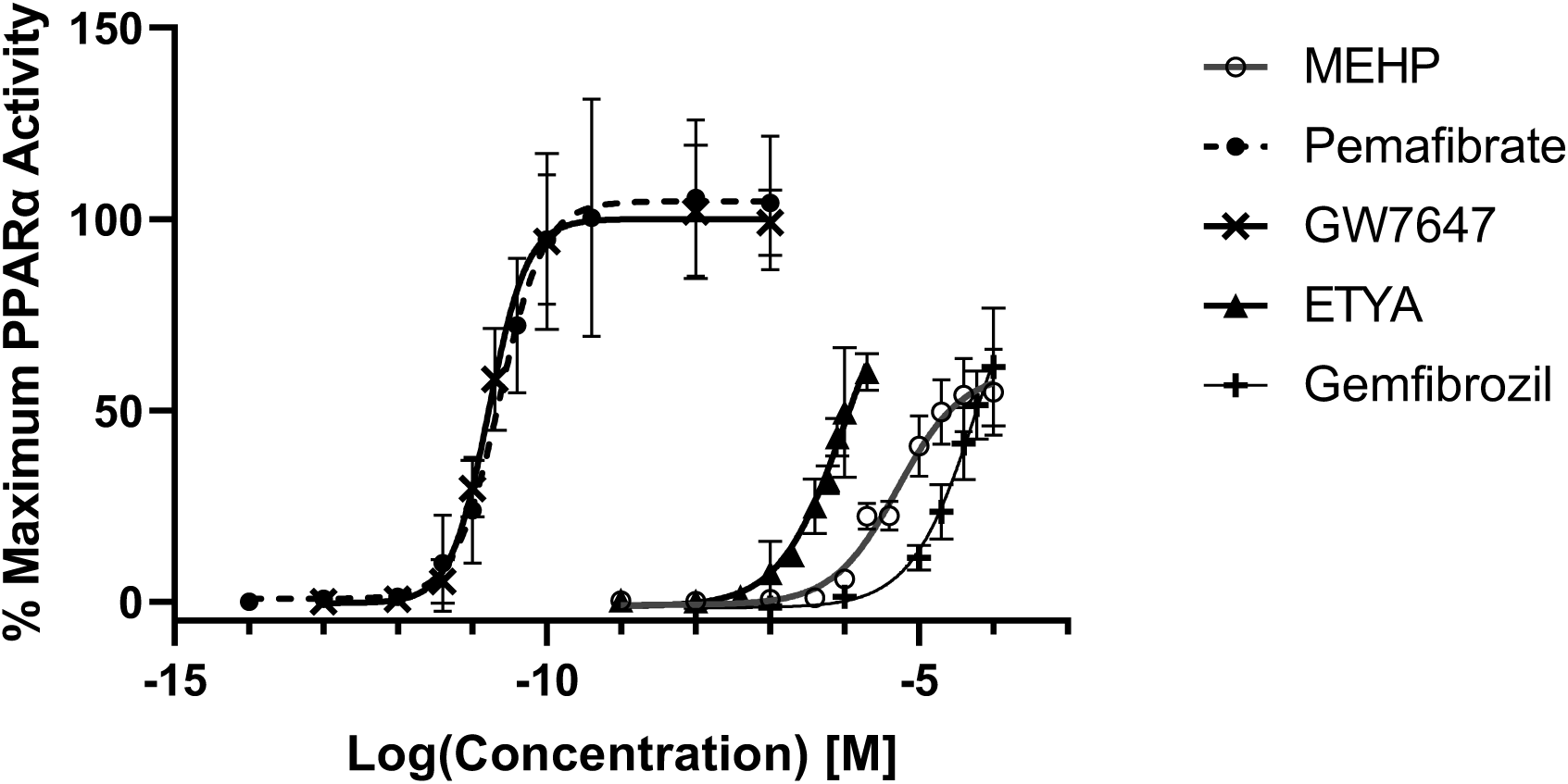
Concentration response curves for activation of human PPARα by established full and partial PPARα agonists. Cos7 cells in 96 well plates were transfected with a full length hPPARα expression construct and a peroxisome proliferator response element-driven luciferase reporter. Cells were treated with vehicle (0.1% DMSO) or varying concentrations of each ligand (GW7647 = 1×10^−12^ to 1×10^−6^ M; pemafibrate = 1×10^−12^ M to 1×10^−7^ M; MEHP = 1×10^−8^ M to 1×10^−4^ M; ETYA = 1×10^−9^ to 2×10^−6^ M; Gemfibrozil 1×10^−8^ to 2×10^−4^ M). Luminescence and fluorescence were measured after 24 hours. Activity was normalized to positive control levels. The lowest concentration is for wells that received Vh only. Curves are fit with a three-parameter Hill function with Hill slope equal to 1. Data points are the mean and standard deviation of four or more independent experiments. Note that the log-scale for concentrations can make maximum effect levels less obvious for some compounds

### 3.2 Mathematical Models Predict hPPARα Activity with Mixtures of Full and Partial Agonists and Antagonists

We used three modeling approaches to predict hPPARα activity for mixtures. Figure 2 shows empirical data (2A) and predicted (2B-D) response surfaces for mixtures of two full hPPARα agonists, GW7647 and pemafibrate. Data are shown in three-dimensional space. The concentration response curve for each chemical alone is visible as the outermost curve in the two prominent planes. Each successive curve is the concentration response curve with a constant concentration of the second full agonist. The response surfaces predicted by GCA and RPF were similar with RMSEs of 11.4 and 11.0, respectively. As expected, ES over-predicted the response for two full agonists, especially at higher concentrations where the effect level predicted by ES was doubled. The high RMSE of 43.8 showed that ES was a poor estimation of the empirical data.

**Figure 2.**
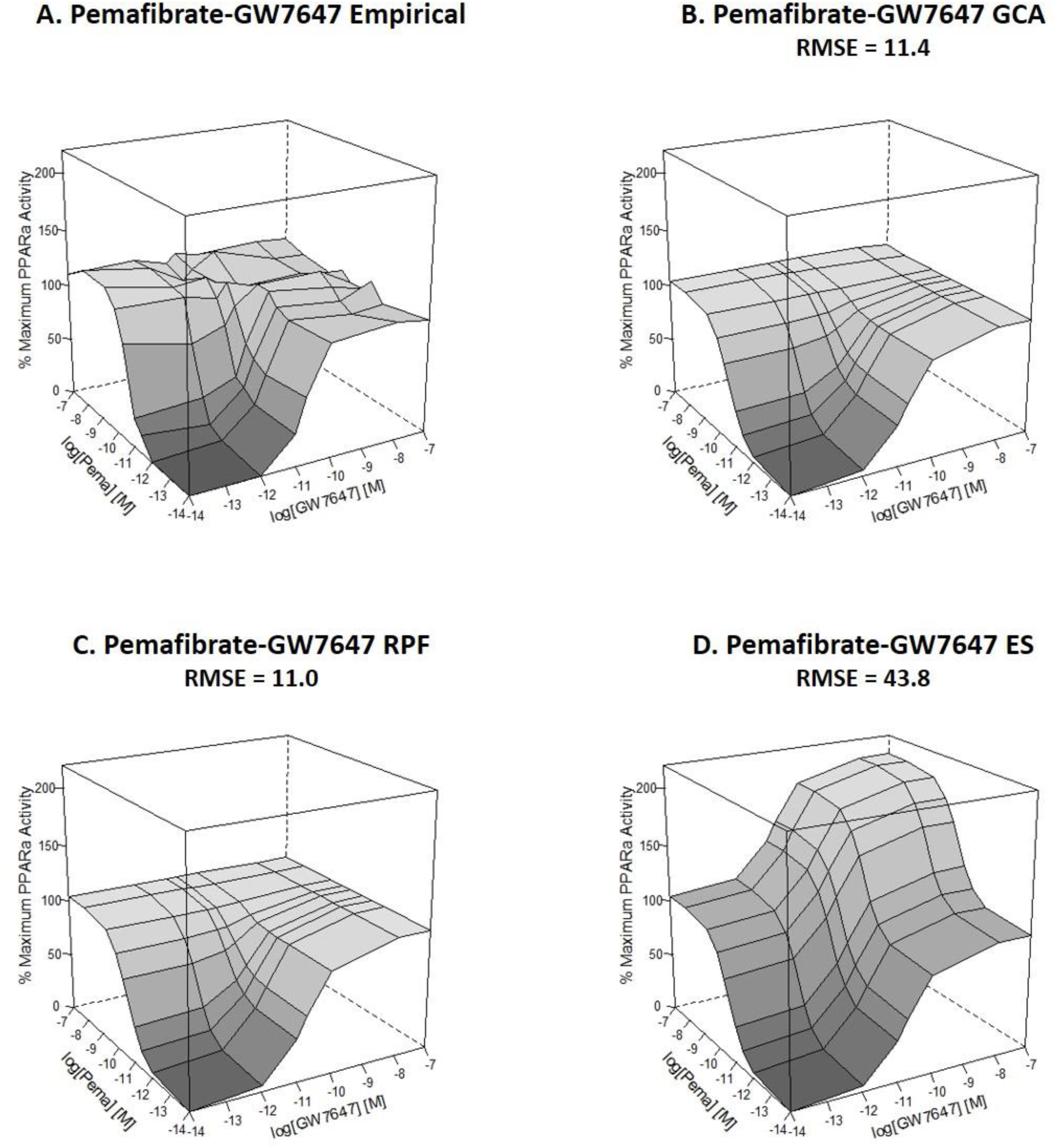
Empirical and predicted response surfaces for mixtures of two full PPARα agonists GW7647 and Pemafibrate (Pema). Cos7 cells in 96 well plates were transfected with a full length hPPARα expression construct and a peroxisome proliferator response element-driven luciferase reporter. Cells were treated in a grid design with eight concentrations of pemafibrate (1×10^−13^ M to 1×10^−7^ M) applied to columns and six concentrations of GW7647 (1×10^−12^M to 1×10^−7^M) applied to rows. Luminescence and fluorescence were measured after 24 hours. Activity was normalized to positive control levels. The lowest concentration is for wells that received Vh only. Data points are the average of four or more independent experiments. A. Empirical data, B. GCA predicted surface, C. RPF predicted surface, D. ES predicted surface.

We hypothesized that GCA, which models hPPARα activity as a function of both potency and efficacy for the mixture components, would more accurately predict empirical data for mixtures of full and partial agonists than RPF, which only accounts for differences in potency between mixture components. Figure 3 shows empirical (3A) and predicted (3B-D) response surfaces for mixtures of the full hPPARα agonist pemafibrate and the partial agonist MEHP. As expected, responses predicted with GCA were more similar to empirical data (RMSE = 8.7) than activity predicted with RPF (RMSE=18.1) and ES (RMSE = 23.8).

**Figure 3.**
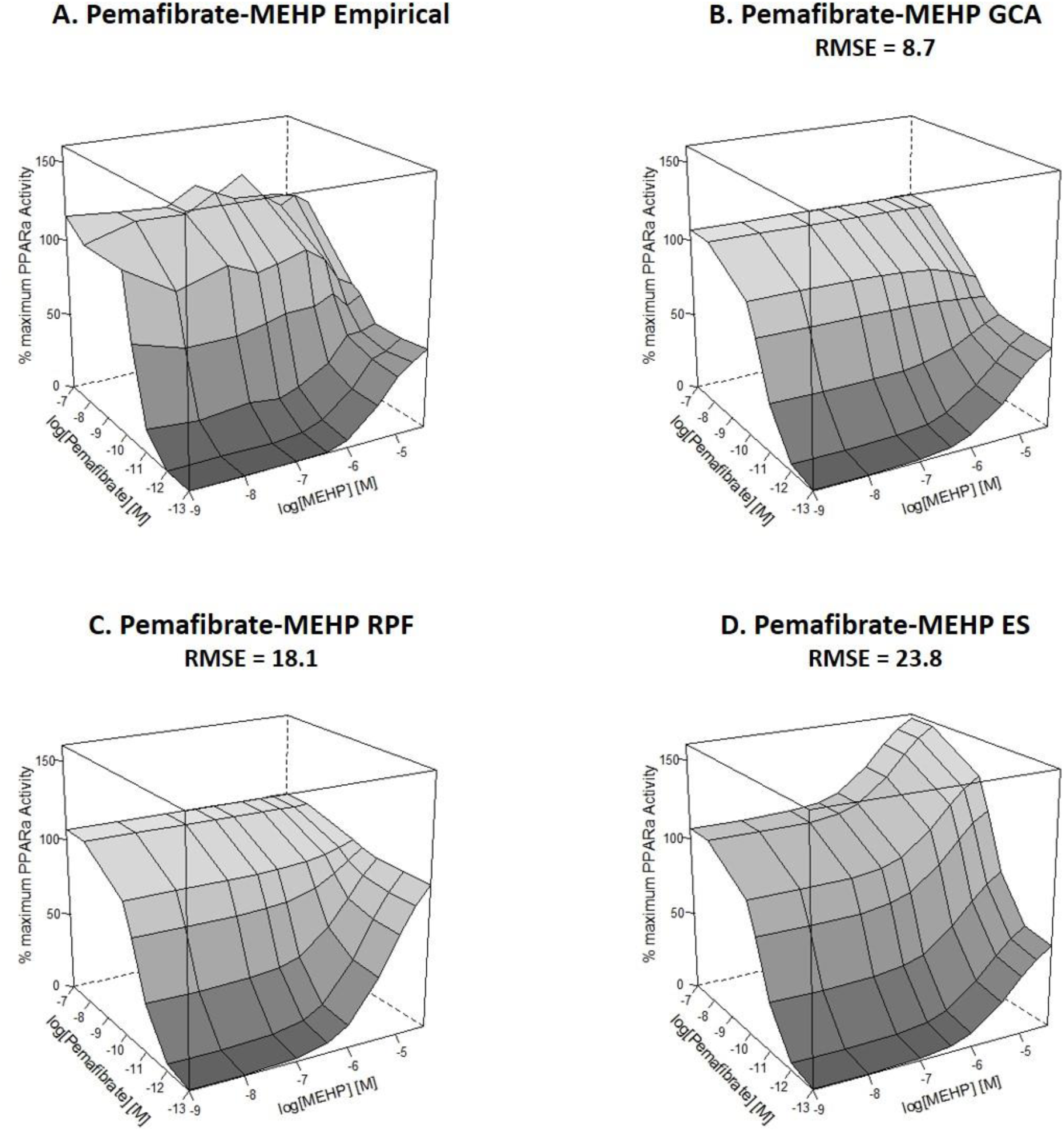
Empirical and predicted response surfaces for mixtures of the full agonist pemafibrate (Pema) with the partial agonist mono(2-diethylhexylphthalate) (MEHP). Cos7 cells in 96 well plates were transfected with a full length hPPARα expression construct and a peroxisome proliferator response element-driven luciferase reporter. Cells were treated in a grid design with six concentrations of pemafibrate (1×10^−12^ M to 1×10^−7^ M) applied to rows and eight concentrations of MEHP (1×10^−8^M to 4×10^−5^M) applied to columns. Luminescence and fluorescence were measured after 24 hours. Activity was normalized to positive control levels. The lowest concentration is for wells that received Vh only. Data points are the average of four or more independent experiments. A. Empirical data, B. GCA predicted surface, C. RPF predicted surface, D. ES predicted surface.

We also tested the ability of GCA to predict activity with mixtures of the full hPPARα agonist pemafibrate and competitive antagonist GW6471. To do so, we first established an inhibition curve with a constant, maximally efficacious concentration of pemafibrate and increasing concentrations of the GW6471 and confirmed that GW6471 was not a hPPARα agonist or inverse agonist (Figure S1). We estimated the equilibrium dissociation constant for GW6471 with a Schild analysis (Schild 1947), as shown in the Supplemental Materials (Tables S3-7, Figures S2-3). The best estimate for the equilibrium dissociation constant for GW6471 is 7.3×10^−9^ M with a 95% confidence interval from 4.5×10^−9^M to 1.0×10^−8^M (Figure S4). GCA can predict activity with mixtures of agonists and competitive antagonists by setting the efficacy for the competitive antagonist to zero and using the equilibrium dissociation constant (Howard and Webster 2009). Empirical and predicted response data are shown in Figure 4. GCA predicted the empirical response data (RMSE=9.6) better than ES (RMSE = 27.2). RPF is not applicable because this method predicts activation of hPPARα by the competitive antagonist.

**Figure 4.**
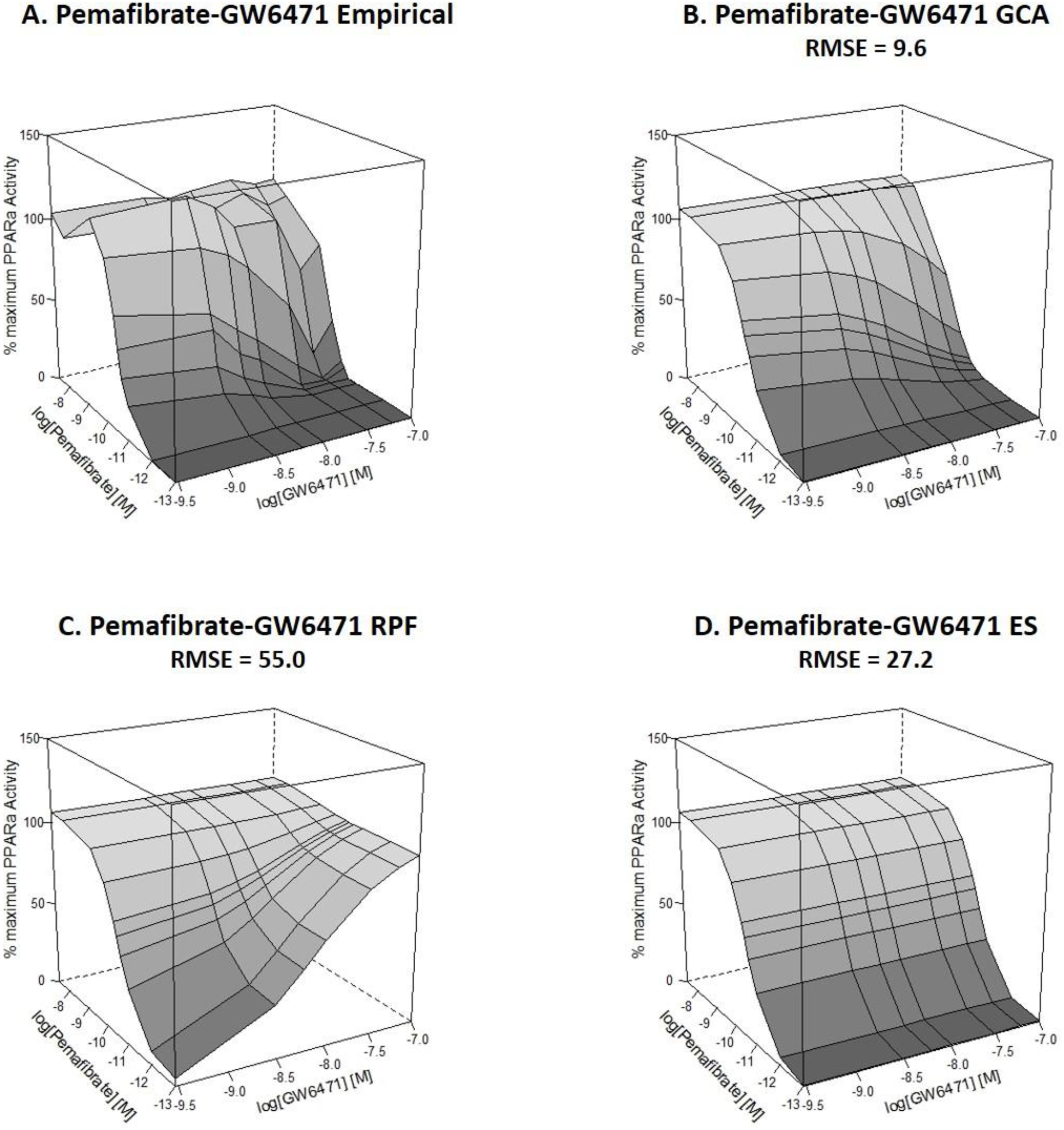
Empirical and predicted response surfaces for mixtures of the full agonist pemafibrate (Pema) with the PPARα competitive antagonist GW6471. Cos7 cells in 96 well plates were transfected with a full length hPPARα expression construct and a peroxisome proliferator response element-driven luciferase reporter. Cells were treated in a grid design with nine concentrations of pemafibrate (1×10^−12^ M to 4×10^−8^ M) applied to columns and six concentrations of GW6471 (3×10^−9^M to 1×10^−7^M) applied to rows. Luminescence and fluorescence were measured after 24 hours. Activity was normalized to positive control levels. The lowest concentration is for wells that received Vh only. Data points are the average of four or more independent experiments. A. Empirical data, B. GCA predicted surface, C. RPF predicted surface, D. ES predicted surface.

### 3.3 PFAS are Full and Partial hPPARα Agonists that Vary in Potency

We generated concentration response curves for PFCAs (PFNA, PFOA, PFHpA, GenX) and PFSAs (PFOS, PFHxS, NBP2) (Figure 5). Concentration response curves are shown on a linear rather than a semi-log scale for two reasons. First, toxicity at high concentrations made it challenging to visualize maximum efficacy on a semi-log scale, which has been noted previously (Buhrke et al. 2013). Second, the closely overlapping concentration responses of PFOA with PFNA and PFOS with NBP2 were poorly distinguished on a semi-log scale. Concentration response curves for the seven PFAS were well fit with three-parameter Hill functions assuming a Hill slope of 1, and the fits were not significantly different from a four parameter model for all PFAS (Table S2).

**Figure 5.**
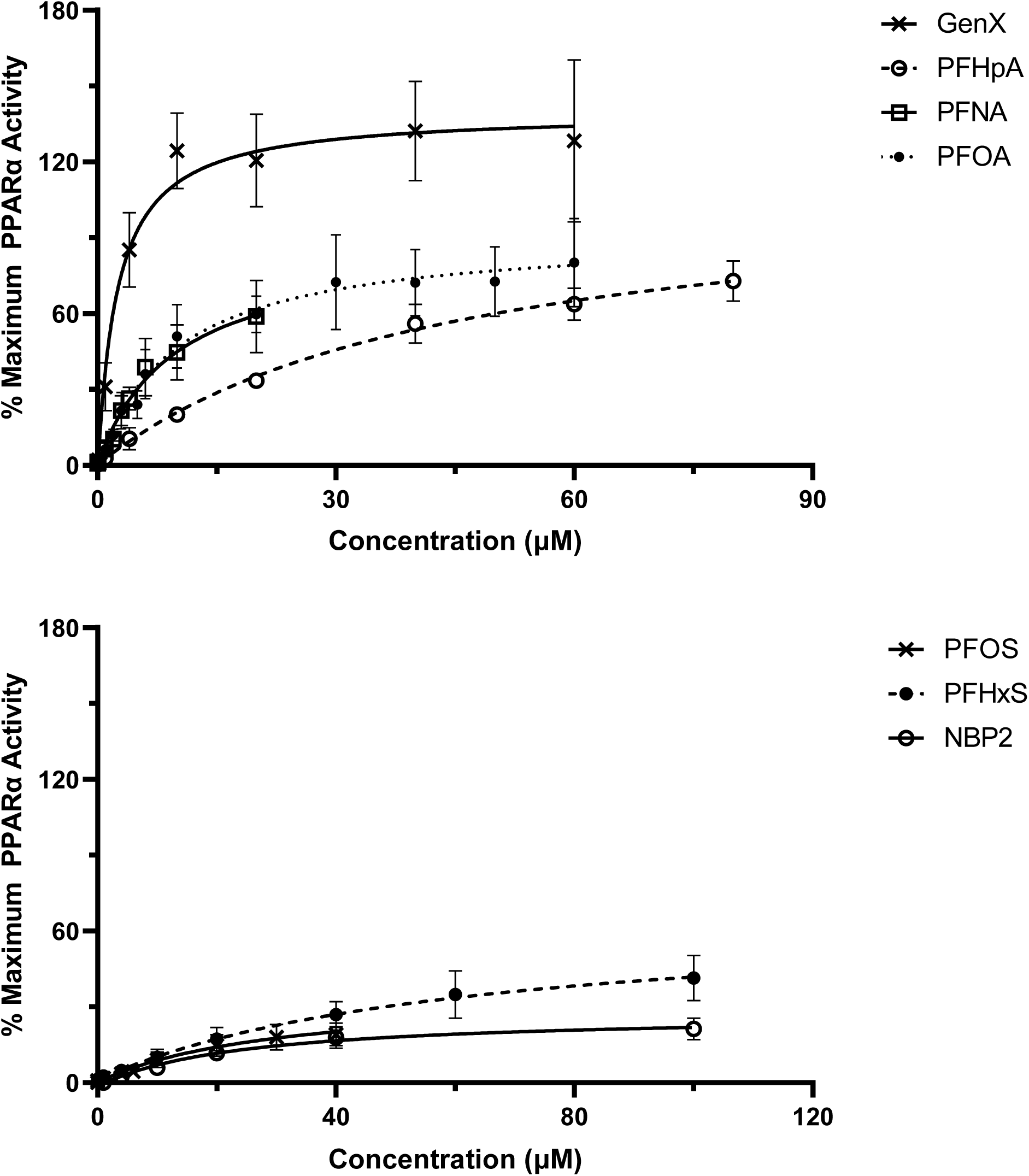
Concentration response curves for A. perfluorinated carboxylic acids and B. perfluorinated sulfonic acids. Cos7 cells in 96 well plates were transfected with a full length hPPARα expression construct and a peroxisome proliferator response element-driven luciferase reporter. Cells were treated with PFAS for 24 hours at varying concentrations (PFOA 4×10^−8^ to 1×10^−4^; PFNA 1×10^−8^ to 1×10^−4^; PFHpA 1×10^−6^ to 4×10^−4^; PFHxS 1×10^−7^ to 2×10^−4^; PFOS 4×10^−8^ to 2×10^−4^; NBP2 1×10^−8^ to 4×10^−4^). Luminescence and fluorescence were measured after 24 hours. Activity was normalized to positive control levels. The lowest concentration is for wells that received Vh only. Curves are fit with a three-parameter Hill function with Hill slope equal to1. Data points are the mean and standard deviation of four or more independent experiments.

PFAS activated hPPARα with differing potency with a ranked order of GenX > PFOA≈PFNA > PFOS≈NBP2 > PFHpA > PFHxS (Table 2). PFCAs acted as full hPPARα agonists with maximum efficacy of 134% for GenX, 113% for PFHpA, 92% for PFOA, and 88% for PFNA compared to positive control activity (Table 2). The three tested PFSA acted as partial hPPARα agonists with maximum efficacies of 67% for PFHxS, 33% for PFOS, and 28% for NBP2 compared with positive control efficacy (Table 2).

**Table 2.** Potency and efficacy with which different PFAS activate full-length human PPARα in Cos7 cells. Parameters estimated from a 3-Parameter Hill Function with Hill slope of 1 in GraphPad Prism v9.0.0.

| PFAS | Potency (EC<sub>50</sub>) $\mu\text{M}$ | Efficacy (% Positive Control<sup>A</sup>) |
| --- | --- | --- |
| PFHpA | 45 | 113 |
| PFOA | 9.5 | 92 |
| PFNA | 9.6 | 88 |
| GenX | 2.1 | 134 |
| PFOS | 24 | 33 |
| PFHxS | 61 | 67 |
| NBP2 | 24 | 28 |

A. Positive control is GW7647 at $1 \times 10^{-6} \text{M}$

### 3.4 Generalized Concentration Accurately Predicts hPPARα Activity of PFAS Mixtures

Because PFCAs acted as full hPPARα agonists and PFSAs acted as partial hPPARα agonists, we hypothesized that 1) GCA would be able to accurately predict the effect of PFCA and PFSA mixtures on hPPARα activity, 2) RPF would overestimate hPPARα activity when a more efficacious compound was used as the reference, and 3) ES would overestimate hPPARα activity at high concentrations. We first tested this hypothesis with two binary mixtures of 1) PFOA and PFOS (Figure 6), and 2) GenX and NBP2 (Figure 7) in grid designs. The mixture surface modeled with GCA was more similar to the empirical mixture surface than predictions by RPF or ES for both the PFOA-PFOS mixture (RMSE 11.9, 19.1, and 19.8, respectively) and for the GenX-NBP2 mixture (RMSE 10.9, 30.0, and 25.7, respectively).

**Figure 6.**
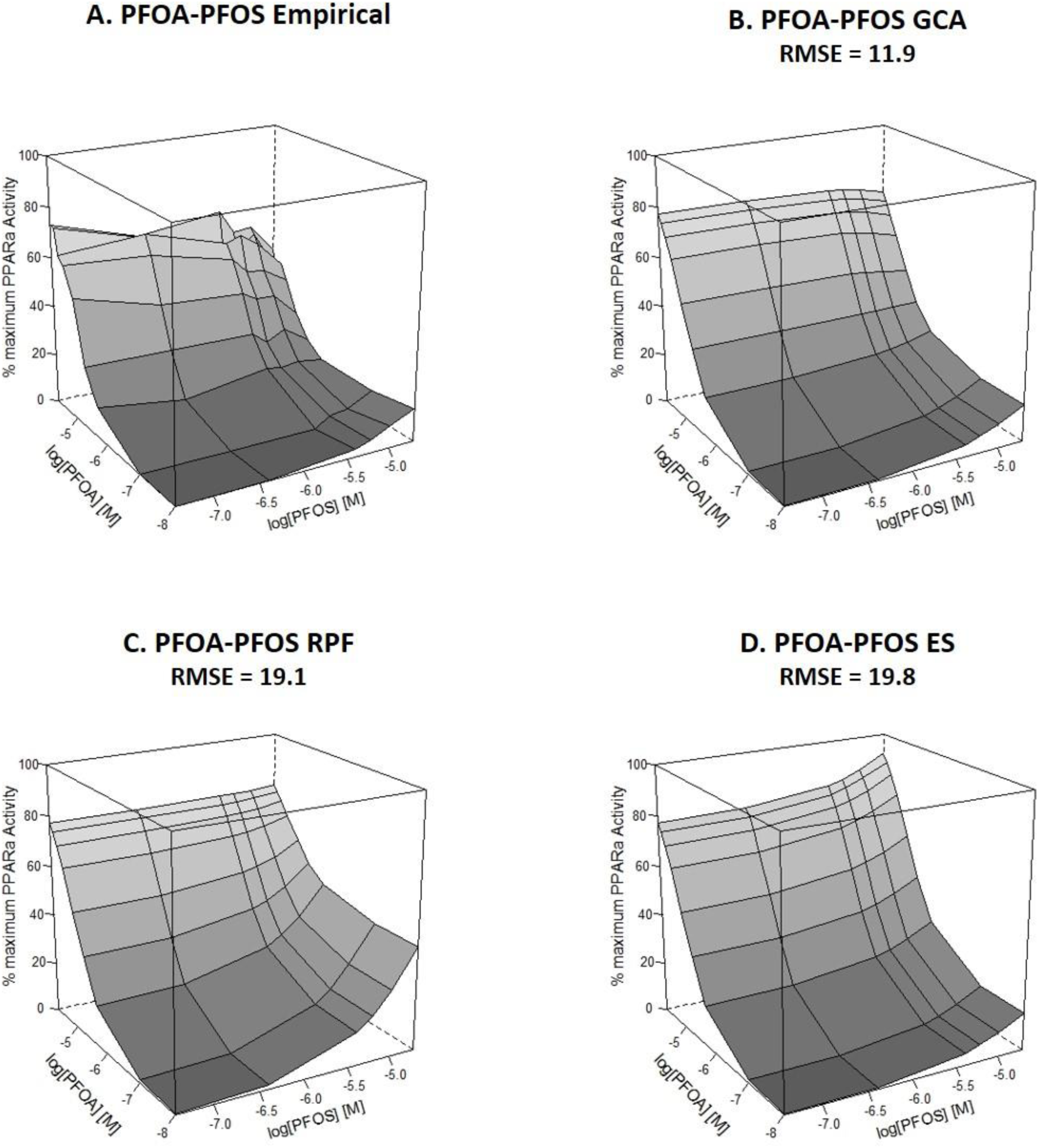
Empirical and predicted concentration response surfaces for mixtures of perfluorooctanoic acid (PFOA) and perfluorooctane sulfonic acid (PFOS). Cos7 cells in 96 well plates were transfected with a full length hPPARα expression construct and a peroxisome proliferator response element-driven luciferase reporter. Cells were treated with increasing PFOA and PFOS concentrations in a grid design. Five concentrations of PFOS were applied to rows (4×10^−7^ to 2×10^−5^ M) and eight concentrations of PFOA were applied to columns (1×10^−7^ to 5×10^−5^ M). Luminescence and fluorescence were measured after 24 hours. Activity was normalized to positive control levels. The lowest concentration is for wells that received Vh only. Data points are the average of four or more independent experiments. A. Empirical data, B. GCA predicted surface, C. RPF predicted surface, D. ES predicted surface.

**Figure 7.**
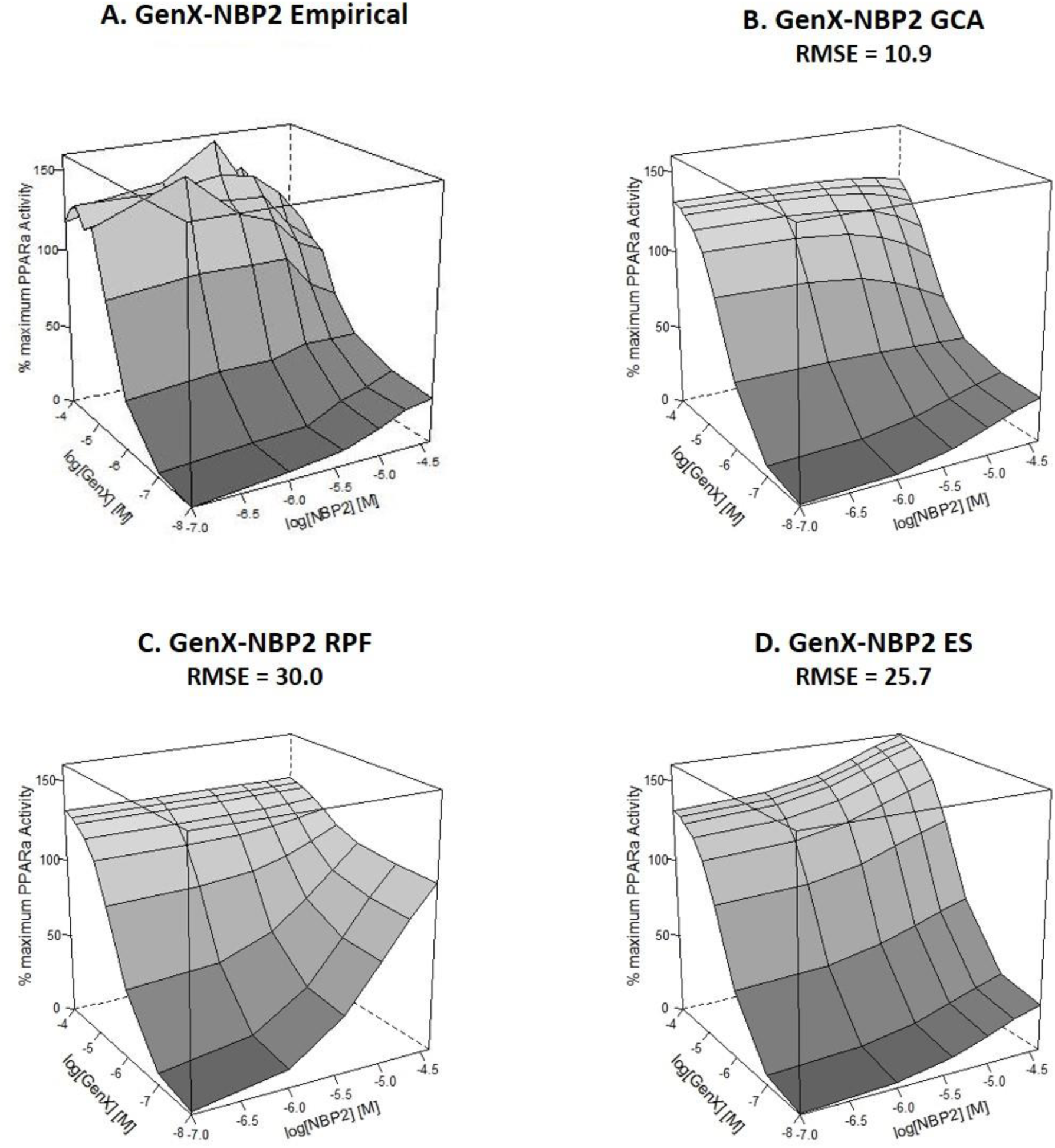
Empirical and predicted concentration response surfaces for mixtures of hexafluoropropylene oxide dimer acid (GenX) and nafion by-product 2 (NBP2). Cos7 cells in 96 well plates were transfected with a full length hPPARα expression construct and a peroxisome proliferator response element-driven luciferase reporter. Cells were treated with increasing GenX and NBP2 concentrations in a grid design. Five concentrations of NBP2 were applied to rows (1×10^−6^ to 4×10^−5^) and eight concentrations of GenX were applied to columns (1×10^−7^ to 1×10^−4^). Luminescence and fluorescence were measured after 24 hours. Activity was normalized to positive control levels. The lowest concentration is for wells that received Vh only. Data points are the average of four or more independent experiments. A. Empirical data, B. GCA predicted surface, C. RPF predicted surface, D. ES predicted surface.

We then assessed the ability of different modeling approaches to predict hPPARα activity with multi-component PFAS mixtures. For a potency-based mixture of the three PFCAs PFHpA, PFOA, and PFNA, both GCA and RPF predicted hPPARα activity within the confidence interval for most observed activity levels (Figure 8A). The RMSE was similar for GCA (8.4) and RPF (11.2) indicating that these two approaches had similar accuracy in predicting the empirical data. ES over-predicted the effects of this PFCA mixture (RMSE = 36.0). GCA more accurately predicted hPPARα activity for equipotent mixtures of the three PFSAs PFHxS, PFOS, and NBP2 with the modeled response falling within the 95% confidence limit for all observed mixture concentrations (Figure 8B). GCA had lower statistical prediction error (RMSE = 3.7) than RPF (RMSE = 23.8) or ES (RMSE = 14.2) for the PFSA mixture. GCA also closely predicted observed hPPARα activity for the mixture with PFOA, PFOS, PFHxS, and PFNA, the four most commonly detected PFAS in the U.S. general population (Figure 8C). The response predicted with GCA was more similar to observed hPPARα activity (RMSE = 5.6) than the response predicted with RPF (RMSE = 23.5) and ES (RMSE = 18.4).

**Figure 8.**
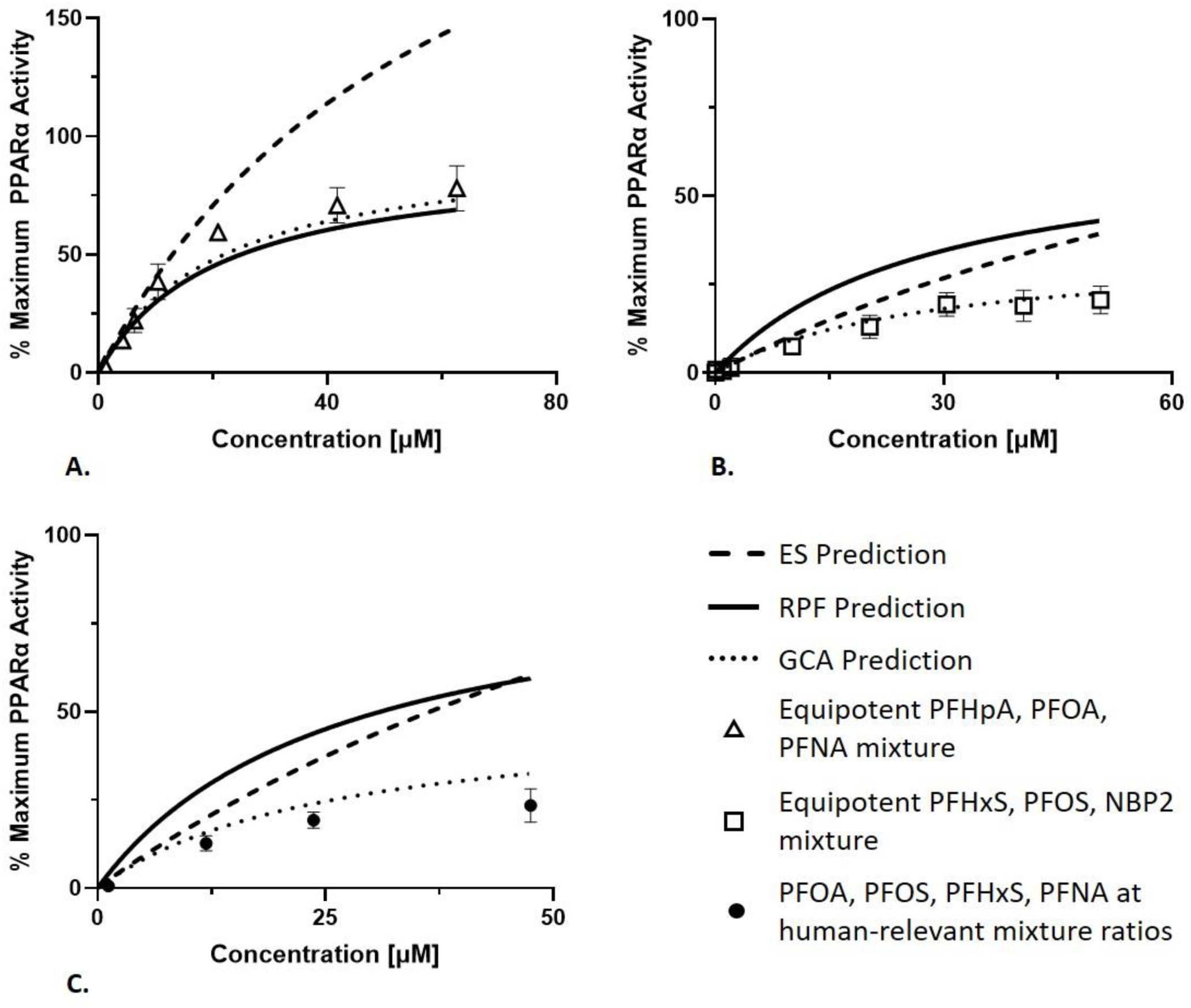
Empirical and modeled PPARα activity for A. Equipotent mixture of PFHpA, PFOA, PFNA, B. Equipotent mixture of PFHxS, PFOS, NBP2, and C. PFOA, PFOS, PFHxS, PFNA at the ratio found in the general U.S. population. Cos7 cells in 96 well plates were transfected with a full length hPPARα expression construct and a peroxisome proliferator response element-driven luciferase reporter. Cells were treated with PFAS mixtures at fixed ratios for 24 hours. The PFCA mixture (A.) and PFSA mixture (B.) are designed so each component contributes equally to the total mixture potency. The PFOA, PFOS, PFHxS, and PFNA mixture has each component at the ratio found in the U.S. general population and is dominated by PFOS. Each mixture was diluted to lower concentrations to measure PPARα activity at different mixture concentrations. Luminescence and fluorescence were measured after 24 hours. Activity was normalized to positive control levels. The lowest concentration is for wells that received Vh only. Data points are the mean and standard deviation of four or more independent experiments.

## 4.0 Discussion

The health effects of chemicals are typically assessed one chemical at a time. However, people are exposed to complex mixtures of environmental chemicals. Here, we evaluated the performance of different mixtures modeling methods (effect summation, relative potency factor and generalized concentration addition) to predict activity of human PPARα (hPPARα) in response to well-characterized synthetic ligands. The PPARα system is of interest due to its critical role in lipid homeostasis and because multiple environmental chemicals such as industrial solvents, plasticizers, and fluorinated surfactants and/or their metabolites interact with this receptor (Kersten 2014; Maloney and Waxman 1999; Shipley et al. 2004; Takacs and Abbott 2007). Because PFAS occur as complex mixtures in the environment, we also tested the accuracy with which modeling approaches predict the effects of PFAS mixtures on hPPARα activity.

### 4.1 Extension of GCA to modeling hPPARα

We began by testing the utility of GCA in predicting the activation of hPPARα by mixtures of well characterized ligands. We found that generalized concentration addition (GCA), a model we developed to accommodate differences in potency and efficacy for mixture components (Howard and Webster 2009), provided accurate predictions of hPPARα activity in Cos7 cells for mixtures of two full hPPARα agonists, mixtures of full and partial agonists, and mixtures of full agonists and antagonists. Potency is a measure of receptor activity expressed as the amount of a ligand that produces a specified effect level, and efficacy is maximum receptor activity induced relative to the maximum attainable receptor activity induced by a positive control ligand. The relative potency factor (RFP) approach, a concentration-additive model that considers only differences in potency for mixture components, provided accurate predictions of hPPARα activity when mixture elements had similar efficacy, but failed to predict activity for mixtures in which individual chemicals differed in efficacy and for mixtures that contained competitive antagonists. Effect summation (ES) over predicted activity of mixtures with full and partial agonists, especially at high concentrations where activity is double counted. ES also failed to predict activity for mixtures of a full agonist and competitive antagonist. These findings are consistent with our previous work showing that GCA accurately predicts activity for mixtures of full and partial agonists for ligand-activated transcription factors with a single binding site (Howard et al. 2010; Watt et al. 2016) and for homodimers in which each monomer binds a ligand independently and then dimerize (Schlezinger et al. 2020a).

### 4.2 Analyses of potency and efficacy of hPPARα activation by PFAS

Our analysis found differences in potency with which PFAS activate hPPARα. We used a full length hPPARα construct, along with a PPARα-driven reporter construct, to characterize the activation of hPPARα by different PFAS. In order of potency, we found that GenX > PFOA ≈ PFNA > PFOS ≈ NBP2 >PFHpA >PFHxS. Our results are largely consistent with findings from hPPARα Gal4 reporter systems in different cell lines, which have shown that PFAS differ in potency with PFCAs being more potent than PFSAs and longer chain PFCAs up to C8/9 being more potent than shorter chain PFCAs (Buhrke et al. 2013; Rosenmai et al. 2018; Wolf et al. 2012; Wolf et al. 2008)(Table S1). In our hands using full length hPPARα, PFOS was more potent than PFHpA, a finding that differs from previous studies using Gal4 reporter systems in which the two PFAS have been tested side by side (Wolf et al. 2012; Wolf et al. 2008).

Differences in potency between PFAS are evident but more nuanced in more complex systems. Studies of endogenous PPARα target gene expression in human-like hepatocytes have found differences in potency when comparing applied PFAS concentrations with PFOA and PFNA more potently inducing most PPARα target genes than PFOS (Louisse et al. 2020). However, in human primary hepatocyte spheroids and *in vivo* with rodent models, differences in the potency with which PFOA and PFOS altered hepatic gene expression and with which PFCAs altered peroxisomal beta oxidation were attenuated with repeated dosing scenarios or when comparing PFAS concentration in the target organ rather than by comparing applied concentration (Kudo et al. 2000; Rowan-Carroll et al. 2021). Thus, future PFAS mixtures modeling experiments should replicate conditions and dosing schemes that are more reflective of chronic exposure scenarios and taking into account pharmacokinetics.

We observed that PFAS activate hPPARα with differing efficacy. The efficacy with which PFCAs activated hPPARα ranged from 88-134% (GenX>PFHpA>PFOA>PFNA) while the efficacy with which PFSAs activated hPPARα ranged from 28-67% for PFSAs (PFHXs>PFOS > NBP2) compared to positive control levels. Effectively, the PFCAs tended to act as full hPPARα agonists while the PFSAs tended to act as partial agonists. Of the 12 studies we found that used reporter systems to define activation of PPARα by PFAS (Table S1), 5 studies suggested the possibility that some PFAS are partial agonists of PPARα (Wolf et al. 2014) or have differences in efficacy (Behr et al. 2020a; Rosenmai et al. 2018; Rosenmai et al. 2016; Vanden Heuvel et al. 2006). Wolf et al., 2014 referenced previous studies to support their conclusion that some PFAS are partial PPARα agonists, rather than highlighting data generated in their study. In HepG2 cells transiently transfected with a PPARα-Gal4 reporter system, Rosenmai et al., 2018 noted that PFHxA, PFHpA, PFOA, PFNA, and PFDA induced higher maximal PPARα activity compared with PFSAs and short and very long chain PFCAs. Behr et al., 2020 also identified differences in “effectiveness” of PPARα transactivation, with GenX being more effective than PFOA in stimulating transcriptional activity by a human PPARα-Gal4 reporter system. None of these studies recognized potential differences in efficacy between PFCA and PFSA. Thus, our finding in Cos7 cells that PFCAs tended to act as full human PPARα agonists and PFSAs tended to act as partial human PPARα agonists is a novel observation.

This analysis differs from most prior reporter assays because we used a full length human PPARα reporter construct rather than a Gal4 reporter construct. In a Gal4 system, the ligand binding domain of PPARα (mouse or human) is fused to the yeast transcription factor Gal4 (Vanden Heuvel et al. 2006; Webster et al. 1988). This construct is structurally distinct from native PPARα, does not require heterodimerization with RXR, the DNA binding partner of PPARα (Gearing et al. 1993; Webster et al. 1988) and binds to an upstream activating sequence, rather than a peroxisome proliferator activated receptor response element, in the reporter gene promoter. Gal4 reporter systems can detect differences in potency and in efficacy (Wilkinson et al. 2008). However, it is unclear how the efficacies in this system translate to a more biologically relevant system. Employing full length reporter constructs that capture both potency and efficacy for the native receptor is an important consideration for modeling the effects of chemical mixtures on key molecular initiating events where individual chemicals differ in both potency and efficacy.

The U.S. Environmental Protection Agency’s ToxCast database screens chemicals, including multiple PFAS, for human PPARα activity using a Gal4 construct driven reporter system in a human hepatocarcinoma cell line. The ToxCast pipeline selects the statistically preferred concentration-response function between three different models including a constant (no-response) function, four parameter Hill model, and a modified four Hill model that accommodates diminished responses at high concentrations to allow for toxicity. As a result, estimates of both potency and efficacy are available through the ToxCast database. In order of potency in this system, PFNA> PFHpA> K-PFHxS> PFOA> PFOS, with potencies ranging from 14.3 μM to 58.9 μM. Efficacies for PFNA, PFHpA, and PFOA were greater than 90% of the maximal activity induced by the positive control. By contrast, PFOS and K-PFHxS induced 66% and 59% of the maximal efficacy induced by the positive control, indicating that PFSAs may act as partial agonists in this system as well, although the relationship between efficacy in Gal4 systems and full length reporter systems is unclear. Importantly, all potency concentrations exceeded the lower bound cytotoxicity limit measured across 35 different cell types, though tested concentrations did not reduce cell viability in the ATG_XTT_Cytotoxicity_up assay in HepG2 cells. Despite this limitation, ToxCast reporter assay data may be amenable to approximate modeling of the effect of complex PFAS mixtures on nuclear receptor activity and validated with further mixtures experiments.

### 4.3 Modeling hPPARα activation by PFAS

The fact that PFCAs and PFSAs have different efficacies for activating hPPARα has significant implications for mixtures modeling. Our results show that GCA more accurately predicts hPPARα activity than RPF and ES for binary mixtures of PFOA and PFOS, mixtures of PFCAs and PFSAs alone, and human-relevant PFAS mixtures. This is not surprising because PFAS concentration response curves do not obey the assumptions for ES or RPF. ES is only applicable for all concentrations when concentration response is linear. For RPF to make accurate predictions, all components must have equal efficacy and have parallel dose response curves (NATO/CCMS Pilot Study on International Information Exchange on Dioxins and Related Compounds 1988; U.S. Environmental Protection Agency 2000). When components of the mixture do not meet these assumptions, the prediction by RFP will be inaccurate, either over- or under-estimating the effect of the mixtures if the referent chemical has a high efficacy or a low efficacy, respectively. Importantly, the GCA prediction is agnostic to the selection of a referent compound because it takes into account the efficacy of each component individually (Howard et al. 2010; Howard and Webster 2009). Therefore, GCA is a significant improvement over other mixtures approaches that have been used historically.

Two prior studies analyzed the effect of PFAS mixtures on mouse PPARα activity in Gal4 reporter systems (Carr et al. 2013; Wolf et al. 2014). Carr et al., 2013 found that all binary combinations of PFOA, PFNA, PFHxS, and PFOS and a human-relevant mixture of these four PFAS resulted in mouse PPARα activity that was less than the activity predicted by a concentration additive model. Our results similarly showed that mixtures of PFOA, PFNA, PFHxS, and PFNA stimulate less activity by hPPARα than would be predicted when applying a simple concentration additive approach such as RPF, which only accounts for differences in PFAS potency. Wolf et al., 2014 found that PPARα activity stimulated by binary combinations PFOA, PFOS, PFHxA, and PFHxS were generally well-predicted by concentration addition and response addition (also called independent action) in lower concentration ranges. But the authors observed both greater and less than additive response at higher concentration levels (Wolf et al. 2014). Our results (Figure 8) showed that ES may fit empirical data in the low concentration regions but does not adequately predict activity at higher concentrations. RPF provides accurate predictions when all mixture components have similar efficacy but can deviate from empirical data, even at low concentrations, when mixture components differ in efficacy. We show that GCA provides more accurate predictions of mixture activity when individual mixture components deviated from the assumptions of RPF and ES. Differences in the ability to predict PFAS mixture activity may be partially attributable to divergent assumptions about the shape of individual concentration response curves: Wolf et al., 2014 applied a concentration additive approach using the average slope across all PFAS in the mixture and did not account for differences in efficacy. Additionally, both studies were conducted in reporter systems with mouse PPARα, which is more responsive to PFAS than the human homolog (Takacs and Abbott 2007; Wolf et al. 2012; Wolf et al. 2008). In the context of similar studies, our results suggest that 1) GCA accurately predicts the effect of PFAS mixtures on hPPARα activity, and 2) RPF could provide conservative estimates of hPPARα activity resulting from PFAS mixture exposure if the reference compound has high efficacy, and underestimate activity if the referent compound has low efficacy.

An RFP-based approach for modeling effects of PFAS mixtures *in vivo* also has been proposed. Bil et al., 2021 derived relative potency factors for 14 PFAS and two PFAS precursors based on liver endpoints including changes in relative and absolute liver weight and liver hypertrophy in male rats following chronic or subchronic oral PFAS exposure. PPARα activation is a critical mechanism contributing to liver hypertrophy in rodents (Hall et al. 2012; Lee et al. 1995). The authors fit dose response curves to for each PFAS that were constrained to have the same slope and efficacy to meet the RPF assumptions. Their results showed that PFCAs and PFSAs with 7-12 perfluorinated carbons have approximately equal or greater potency than PFOA while shorter and longer chain PFAS, carboxylic acid ethers (GenX), and fluorotelomer alcohols have lower potency. PFOS, PFNA, PFUnDA, and PFDoDa all had RPFs greater than one indicating they are more potent than PFOA in this system. While this analysis is an important step forward for assessing risk associated with exposure to PFAS mixtures, the authors did not evaluate the goodness of fit for the assumptions of equal slope and efficacy for the underlying dose response curves for individual PFAS, which may limit the accuracy of RPF predictions. If differences in efficacy are evident, *in vivo* RPFs may over or under predict the effect of PFAS mixtures in liver endpoints, depending on the reference compound. Further, differences in potency may also reflect differences in distribution and elimination between PFAS in both a single species and across species. This has been suggested by another group that compared the potency ranking of PFOA, PFHxA, PFBA, and GenX for effects on increased liver weight for oral administered dose and modeled serum and target organ PFAS concentrations (Gomis et al. 2018). Their analysis found that differences in potency were reduced when comparing modeled liver PFAS concentrations rather than oral or serum measures (Gomis et al. 2018). Human relevant systems remain essential to validate the effects of PFAS mixtures on important molecular initiating events.

Studies of the health effects of PFAS mixtures in human-relevant systems remains an important data gap. Our analysis provides an essential contribution by assessing the effects of PFAS mixtures on a human-relevant molecular initiating event in a simplified system. An important limitation remains the interactive effects of PFAS on downstream health effects given the ability of PFAS to interact with multiple nuclear receptors and other molecular initiating events. Future analyses should expand this work on the effects of PFAS mixtures on human relevant molecular initiating events by examining mixtures effects in intact cells where multiple molecular initiating events contribute to proximate effects and functional changes.

## 5.0 Conclusions

We used a simplified system with Cos7 cells transiently transfected with full length hPPARα to test the ability of modeling approaches that can accommodate diverse PPARα ligands to predict the effect of PFAS mixtures on hPPARα activity. In this system, PFCAs acted as full and PFSAs acted as partial human PPARα agonists that differ in potency. We showed that hPPARα activity induced by PFAS mixtures can be accurately predicted by GCA while RPF predicted activity when mixture components had similar efficacy but did not when components differed in efficacy. ES predictions were accurate at lower concentrations and overestimated PPARα activity at higher concentrations. Our results are consistent with prior findings that concentration additive approaches that consider only differences in chemical potency may over predict the effects of PFAS mixtures on PPARα activity. This analysis begins to fill an important data gap by showing that GCA can predict the effect of PFAS mixtures on a human-relevant molecular initiating event. Models that can predict the effects of PFAS mixtures on molecular initiating events and downstream health effects are a critical tool to assess health risks from mixtures of PFAS in the environment.

## Supporting information

PFAS Mixtures Supplemental Material

## 6.0 Acknowledgements

The authors thank Nathan Burritt and Hannah Puckett for their technical support, patience, and guidance during data collection for this manuscript.

