## Supplementary material for "Predicting the Effects of Per- and Polyfluoroalkyl Substance Mixtures on Peroxisome Proliferator-Activated Receptor Alpha Activity *in Vitro*": PFAS Mixtures Supplemental Material

**<u>Table of Contents</u>**

| | Page |
|---|---|
| **Table S1.** Reporter-based analyses of PPARα activation by PFAS. | **2** |
| **Table S2**. Model fits for full and partial PPARα agonists and PFAS | **5** |
| **Table S3.** Parameters for individual dose response curves for pemafibrate with increasing concentrations of the PPARα antagonist GW6471 | **7** |
| **Table S4**. Parameters for pemafibrate dose response curves with increasing concentrations of GW6471 constrained to mean values from Supplemental Table 2 | **7** |
| **Table S5.** Input values and calculated AICs for individual and mean fits | **8** |
| **Table S6.** Dose ratio values using mean curve fits | **8** |
| **Table S7.** Schild regression goodness of fit, confidence interval, and equation | **8** |
| **Figure S1**. GW6471 inhibition curve for pemafibrate. | **9** |
| **Figure S2**. Unconstrained pemafibrate dose response curves with increasing concentrations of GW6471 | **10** |
| **Figure S3**. Constrained pemafibrate dose response curves with the indicated dose of GW6471 | **11** |
| **Figure S4**. Schild regression to estimate equilibrium dissociation constant of GW6471 | **12** |
| **References** | **13** |

**Table S1. Reporter-based analyses of PPARα activation by PFAS.**

| Species | Construct Type | Serum | Positive Control | | PFAS | | | Ref |
|---|---|---|---|---|---|---|---|---|
| | | | µM | Max Activity (Fold Increase) | | µM | Max Activity (Fold Increase) | |
| **Mouse** | Full length | No | Wy – 20 | 16.2 | PFOA | EC$_{90}$ - 10 | 17.2 | [1] |
| **Human** | | | Wy – 20 | 5.7 | PFOA | EC$_{90}$ - 20 | 8.9 | |
| **Mouse** | Full length | No | Wy – 20 | 4 | *PFOS* | *EC$_{90}$ - 32* | *4.6* | [2] |
| **Human** | | | Wy – 20 | 4 | *PFOS* | *EC$_{90}$ - 84* | *3.4* | |
| **Mouse** | | | Wy – 20 | 4 | *FOSA* | *EC$_{90}$ - 29* | *3.9* | |
| **Human** | | | Wy – 20 | 4 | *FOSA* | *EC$_{90}$ - 41* | *3.0* | |
| **Mouse** | Gal4 | Yes | Cipro[a] – 9.5 (EC$_{50}$) | 17.8 | lin-PFOA | EC$_{50}$ – 60.3 | 9.3 | [3] |
| **Human** | | | Cipro[a] – 8.6 (EC$_{50}$) | 13 | lin-PFOA | EC$_{50}$ – 45.2 | 10.2 | |
| **Mouse** | Gal4 | No | Wy – 30 | 23 | PFOA | 40[b] | 2.5 | [4] |
| **Human** | | | Wy – 50 | 3 | PFOA | 40 | 1.8 | |
| **Mouse** | | | Wy – 30 | 23 | *PFOS* | *120* | *1.5* | |
| **Human** | | | Wy – 50 | 3 | *PFOS* | *250* | *No activity* | |
| **Mouse** | Gal4 | No | Wy – 10 | Not reported | PFBA | EC$_{20}$ – 51[c] | 0.45[d] | [5, 6] |
| **Human** | | | | | PFBA | EC$_{20}$ –75 | 0.30 | |
| **Mouse** | | | | | PFHxA | EC$_{20}$ –38 | 0.5 | |
| **Human** | | | | | PFHxA | EC$_{20}$ –47 | 0.5 | |
| **Mouse** | | | | | PFOA | EC$_{20}$ –6 | 1 | |
| **Human** | | | | | PFOA | EC$_{20}$ –16 | 0.5 | |
| **Mouse** | | | | | PFNA | EC$_{20}$ –5 | 1.25 | |
| **Human** | | | | | PFNA | EC$_{20}$ –11 | 0.5 | |
| **Mouse** | | | | | PFDA | EC$_{20}$ –20 | 1 | |
| **Human** | | | | | PFDA | No activity | | |
| **Mouse** | | | | | *PFBS* | *EC$_{20}$ – 317* | *0.15* | |
| **Human** | | | | | *PFBS* | *EC$_{20}$ – 206* | *0.25* | |
| **Mouse** | | | | | *PFHxS* | *EC$_{20}$ – 76* | *0.3* | |
| **Human** | | | | | *PFHxS* | *EC$_{20}$ – 81* | *0.25* | |
| **Mouse** | | | | | *PFOS* | *EC$_{20}$ – 94* | *0.55* | |
| **Human** | | | | | *PFOS* | *EC$_{20}$ – 262* | *0.25* | |
| **Mouse** | Gal4 | Not indicated | Wy – 10 | Not reported | PFPeA | EC$_{20}$ – 45[c] | 0.5[d] | [7, 8] |
| **Human** | | | | | PFPeA | EC$_{20}$ – 52 | 0.4 | |
| **Mouse** | | | | | PFHpA | EC$_{20}$ – 11 | 0.6 | |
| **Human** | | | | | PFHpA | EC$_{20}$ – 15 | 0.4 | |

| | | | | | | | | |
|---|---|---|---|---|---|---|---|---|
| **Mouse** | | | | | PFOA | $EC_{20}$ – 7 | 1 | |
| **Human** | | | | | PFOA | $EC_{20}$ – 7 | 0.9 | |
| **Mouse** | | | | | PFUnDA | $EC_{20}$ – 8 | 1.1 | |
| **Human** | | | | | PFUnDA | $EC_{20}$ – 86 | 0.3 | |
| **Mouse** | | | | | PFDoDA | $EC_{20}$ – 33 | 0.6 | |
| **Human** | | | | | PFDoDA | No activity | | |
| **Human** | Gal4 | Not indicated | GW7647 - 1 | 5 | PFBA | $EC_{20}$ – 27.4 | Not reported | [9] |
| | | | | | PFHxA | $EC_{20}$ – 12.2 | | |
| | | | | | PFHpA | $EC_{20}$ – 5.3 | | |
| | | | | | PFOA | $EC_{20}$ – 0.9 | | |
| | | | | | PFNA | $EC_{20}$ – 21.3 | | |
| | | | | | PFDA | $EC_{20}$ – 28.4 | | |
| | | | | | PFDoDA | $EC_{20}$ – >20 | | |
| **Human** | Gal4 | Not indicated | WY (concentration not reported) | Not reported | PFBA | 30[e] | 1.8 | [10, 11] |
| | | | | | PFPeA | 30 | 1.8 | |
| | | | | | PFHxA | 100 | 2.5 | |
| | | | | | PFHpA | 30 | 2.1 | |
| | | | | | PFOA | 30 | 2.5 | |
| | | | | | PFNA | 100 | 2.5 | |
| | | | | | PFDA | 100 | 2.5 | |
| | | | | | PFUnDA | No Activity | | |
| | | | | | PFDoDa | 100 | 1.5 | |
| | | | | | PFTeDA | 10 | 2 | |
| | | | | | *PFBS* | *30* | *1.5* | |
| | | | | | *PFHxS* | *30* | *1.8* | |
| | | | | | *PFOS* | *No Activity* | | |
| | | | | | *FOSA* | *No Activity* | | |
| **Human** | Gal4 | Not indicated | GW7647 - 1 | 16 | PFBA | 50[f] | 2 | [12] |
| | | | | | PFHxA | 50 | 2 | |
| | | | | | PFOA | 50 | 2.5 | |
| | | | | | *PFBS* | *No Activity* | | |
| | | | | | *PFHxS* | *100* | *2* | |
| | | | | | *PFOS* | *100* | *2* | |
| | | | | | PMPP | 50 | 2 | |
| | | | | | GenX | 25 | 5 | |

Note: When numerical data were not reported, activity was estimated from the plots.

a – The $EC_{50}$ is reported because the concentration that induced the maximal activity was not reported.

b – No potency comparison was reported. The concentration listed is that which induced maximal activity.

c – The $EC_{20}$ is reported because the concentration that induced the maximal activity was not reported.

d – Units are log relative fluorescence units.

e – Only 3 concentrations were analyzed (10, 30 and 100 µM). The lowest concentration that significantly induced activity is provided in the table, along with the activity at that concentration.

f – Only 3 concentrations were analyzed (25, 50 and 100 µM). The lowest concentration that significantly induced activity is provided in the table, along with the activity at that concentration.

Abbreviations: Ammonium perfluoro(2-methyl-3-oxahexanoate) (GenX), ciprofibrate (Cipro), per-fluorooctanesulfonamide (FOSA), perfluorobutanoic acid (PFBA), perfluorobutane sulfonate (PFBS), perfluorodecanoic acid (PFDA), perfluorododecanoic acid (PFDoDA), perfluoroheptanoic acid (PFHpA), perfluorohexanoate (PFHxA), perfluorohexane sulfonate (PFHxS), perfluorooctanoic acid (PFOA), perfluorooctane sulfonate (PFOS), perfluorononanoic acid (PFNA), perfluoropentanoic acid (PFPeA), perfluorotetradecanoate (PFTeDA), perfluoroundecanoic acid (PFUnDA), 3H-perfluoro-3-[(3-methoxypropoxy) propanoic acid] (PMPP) and WY - WY-14643.

**Table S2**. Model fits for full and partial PPARα ligands and PFAS compared in GraphPad Prism v 9.0.0 with the "log(agonist) vs. response (three parameters)" model versus the log(agonist) vs. response—Variable slope (four parameters) model. P-values tests whether the statistical fit is significantly better with a four- or three-parameter Hill function.

| | **3 Parameter** | **4 Parameter** | **P-Value** |
|---|---|---|---|
| GW7647 | AIC = 648 | AIC = 648 | *0.22* |
| EC50 | 1.8E-11 | 1.8E-11 | |
| Maximum | 99 | 97 | |
| Hill Slope | 1 | 1.3 | |
| PEMAFIBRATE | AIC = 1046 | AIC = 1047 | *0.36* |
| EC50 | 2.2E-11 | 2.3E-11 | |
| Maximum | 104 | 103 | |
| Hill Slope | 1 | 1.2 | |
| GEMFIBROZIL | AIC = 132 | AIC = 134 | *0.42* |
| EC50 | 5.8E-05 | 3.6E-05 | |
| Maximum | 98 | 78 | |
| Hill Slope | 1 | 1.3 | |
| ETYA | AIC = 377 | AIC = 377 | *0.12* |
| EC50 | 9.9E-07 | 6.5E-07 | |
| Maximum | 94 | 74 | |
| Hill Slope | 1 | 1.4 | |
| Mono(2-ethylhexyl) phthalate (MEHP) | AIC = 301 | AIC = 301 | *0.12* |
| EC50 | 5.2E-06 | 4.80E-06 | |
| Maximum | 60.1 | 57 | |
| Hill Slope | 1 | 1.2 | |
| Perfluorooctanoic acid (PFOA) | AIC = 594 | AIC = 593 | *0.08* |
| EC50 | 9.5E-06 | 7.6E-06 | |
| Maximum | 92 | 81 | |
| Hill Slope | 1 | 1.3 | |
| Perfluorononanoic acid (PFNA) | AIC = 330 | AIC = 330 | *0.14* |
| EC50 | 9.6E-06 | 6.0E-06 | |
| Maximum | 88 | 69 | |
| Hill Slope | 1 | 1.3 | |
| Perfluoroheptanoic acid (PFHpA) | AIC = 95 | AIC = 97 | *0.72* |
| EC50 | 4.5E-05 | 3.7E-05 | |
| Maximum | 113 | 104 | |
| Hill Slope | 1 | 1.1 | |
| Perfluorohexane sulfonic acid (PFHxS) | AIC = 210 | AIC = 212 | *0.74* |
| EC50 | 6.1E-05 | 5.1E-05 | |
| Maximum | 67 | 61 | |
| Hill Slope | 1 | 1.1 | |
| Perfluorooctane sulfonic acid (PFOS) | AIC = 133 | AIC = 134 | *0.27* |

| | | | |
|---|---|---|---|
| EC50 | 2.4E-05 | 1.4E-05 | |
| Maximum | 33 | 24 | |
| Hill Slope | 1 | 1.4 | |
| Nafion by-product 2 (NBP2) | AIC = 62 | AIC = 63 | 0.32 |
| EC50 | 2.4E-05 | 1.9E-05 | |
| Maximum | 28 | 24 | |
| Hill Slope | 1 | 1.3 | |
| Perfluoro(2-methyl-3-oxahexanoic) acid (GenX) | AIC =218 | AIC =217.1 | 0.08 |
| EC50 | 2.1E-06 | 2.3E-06 | |
| Maximum | 134 | 128 | |
| Hill Slope | 1 | 1.6 | |

EC50=half-maximal effective concentration; AIC= Akaike information criterion

**Table S3**. Comparing potency and efficacy for human PPARa activity in cos-7 cells transiently transfected with full-length human PPARα compared with HepG2 cells transfected with a human PPARa Gal4 construct.

| | **Potency (µM)** | | **Efficacy[1]** | |
|---|---|---|---|---|
| | **ToxCast (HepG2)** | **Cos-7** | **ToxCast (HepG2)** | **Cos-7** |
| **PFNA** | 14.3 | 9.6 | 123 | 88 |
| **PFHpA** | 15.9 | 45 | 95 | 113 |
| **PFHxS[2]** | 17.1 | 61 | 59 | 67 |
| **PFOA** | 21.8 | 9.6 | 132 | 92 |
| **PFOS** | 58.9 | 24 | 66 | 33 |

1. Efficacy is the percent of maximal activity induced by the positive control chemical (GW0742 for HepG2, GW7647 for Cos-7)

**Table S4.** Parameters and fits for individual dose response curves for pemafibrate with increasing concentrations of the PPARα antagonist GW6471.

| | **No GW6471** | **6.00x10$^{-9}$** | **1.00x10$^{-8}$** | **3.00x10$^{-8}$** | **6.00x10$^{-8}$** | **1.00x10$^{-7}$** |
|---|---|---|---|---|---|---|
| **Bottom** | -5.579 | -5.954 | -6.654 | -8.046 | -4.889 | -4.366 |
| **Top** | 108.3 | 109 | 103.2 | 109.6 | 100.1 | 99.61 |
| **LogEC50** | -10.57 | -10.3 | -10.21 | -9.877 | -9.479 | -9.373 |
| **HillSlope** | 1 | 1 | 1 | 1 | 1 | 1 |
| **EC50** | 2.714x10$^{-11}$ | 4.972x10$^{-11}$ | 6.106x10$^{-11}$ | 1.33x10$^{-10}$ | 3.32x10$^{-10}$ | 4.24x10$^{-10}$ |
| Span | 113.9 | 114.9 | 109.9 | 117.7 | 105 | 104 |
| **Sum of Squares** | 15342 | 6571 | 3448 | 7889 | 2805 | 2253 |
| **AICc** | 429 | 213.2 | 187.4 | 220.5 | 179.2 | 161.3 |
| **# Data Points** | 80 | 40 | 40 | 40 | 40 | 37 |

**Table S5**. Parameters and model fits for dose response curves constrained to the mean maximum value from curves in Supplemental Figure 2.

| | **No GW6471** | **6.00x10$^{-9}$** | **1.00x10$^{-8}$** | **3.00x10$^{-8}$** | **6.00x10$^{-8}$** | **1.00x10$^{-7}$** |
|---|---|---|---|---|---|---|
| **Bottom** | -5.704 | -6.07 | -6.535 | -8.191 | -4.642 | -4.103 |
| **Top** | = 105.6 | = 105.6 | = 105.6 | = 105.6 | = 105.6 | = 105.6 |
| **LogEC50** | -10.59 | -10.33 | -10.2 | -9.905 | -9.433 | -9.321 |
| **HillSlope** | 1 | 1 | 1 | 1 | 1 | 1 |
| **EC50** | 2.56x10$^{-11}$ | 4.678x10$^{-11}$ | 6.379x10$^{-11}$ | 1.24x10$^{-10}$ | 3.69 x10$^{-10}$ | 4.77 x10$^{-10}$ |
| **Span** | = 111.3 | = 111.6 | = 112.1 | = 113.8 | = 110.2 | = 109.7 |
| **Sum of Squares** | 15518 | 6696 | 3504 | 8033 | 3023 | 2469 |
| **AICc** | 427.7 | 211.5 | 185.6 | 218.8 | 179.7 | 162.2 |
| **# Y values** | 80 | 40 | 40 | 40 | 40 | 37 |

**Table S6.** Input values and calculated AICs for individual and mean fits.

| | Sum of Squares | Number of Observations | Number of Groups | AIC |
|---|---|---|---|---|
| **Individual Fits** | 44810 | 317 | 14 | 1599 |
| **Mean Fits** | 45893 | 317 | 8 | 1594 |

**Table S7.** Dose ratio values using mean curve fits.

| | $6.00x10^{-9}$ | $1.00x10^{-8}$ | $3.00x10^{-8}$ | $6.00x10^{-8}$ | $1.00x10^{-7}$ |
|---|---|---|---|---|---|
| **DR** | 1.8 | 2.5 | 4.8 | 14.4 | 18.6 |
| **DR-1** | 0.8 | 1.5 | 3.8 | 13.4 | 17.6 |
| **Log(DR-1)** | -0.084 | 0.173 | 0.585 | 1.127 | 1.246 |

**Table S8.** Schild regression goodness of fit, confidence interval, and equation. Parameters were calculated in GraphPad Prism version 9.0.0 with a simple linear regression equation.

| | Log(DR-1) | Unity |
|---|---|---|
| **Best-fit values** | | |
| **Slope** | 1.119 | 1 |
| **Y-intercept** | 9.109 | 8.14 |
| **X-intercept** | -8.138 | -8.138 |
| **1/slope** | 0.8934 | 0.9997 |
| **Std. Error** | | |
| **Slope** | 0.0836 | 0.07471 |
| **Y-intercept** | 0.636 | 0.5684 |
| **95% Confidence Intervals** | | |
| **Slope** | 0.8533 to 1.385 | 0.7626 to 1.238 |
| **Y-intercept** | 7.085 to 11.13 | 6.332 to 9.949 |
| **X-intercept** | -8.348 to -7.993 | -8.348 to -7.993 |
| **Goodness of Fit** | | |
| **R squared** | 0.9835 | 0.9835 |
| **Sy.x** | 0.08589 | 0.07675 |
| **Is slope significantly non-zero?** | | |
| **F** | 179.3 | 179.3 |
| **DFn, DFd** | 1, 3 | 1, 3 |
| **P value** | 0.0009 | 0.0009 |
| **Equation** | Y = 1.119*X + 9.109 | Y = 1.000*X + 8.140 |

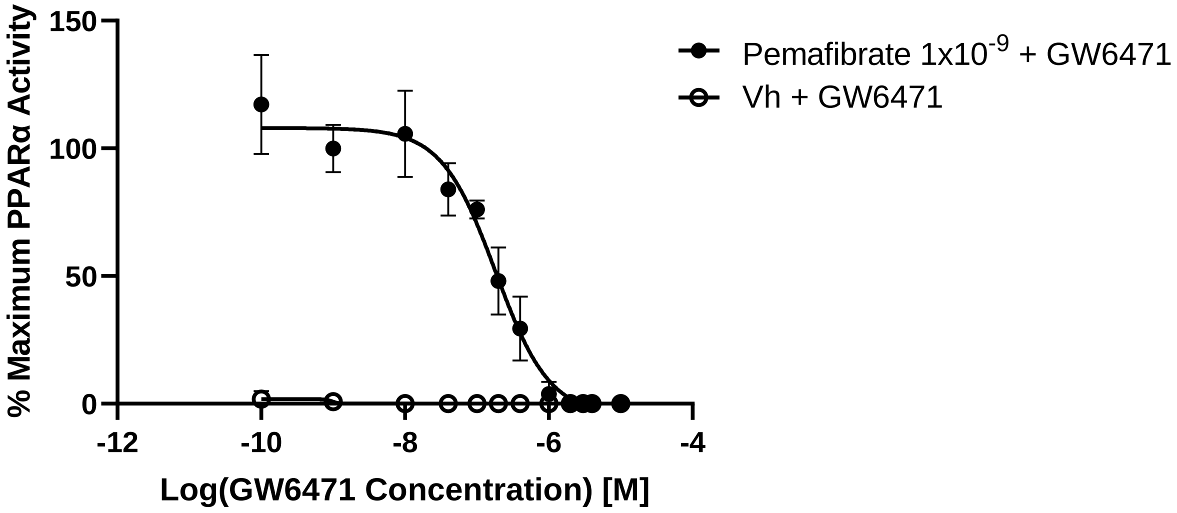

**Figure S1**. GW6471 inhibition curve for pemafibrate. Cos-7 cells were plated in serum free medium and dosed with 1) a maximally efficacious concentration of pemafibrate ($1x10^{-9}$ M) and increasing concentrations of the PPARα antagonist GW6471 or 2) increasing GW6471 concentrations with vehicle (DMSO). Luminescence and fluorescence were measured after 24 hrs. Data were calculated as described in the Methods. Data points show the mean and standard deviation of 4 independent experiments.

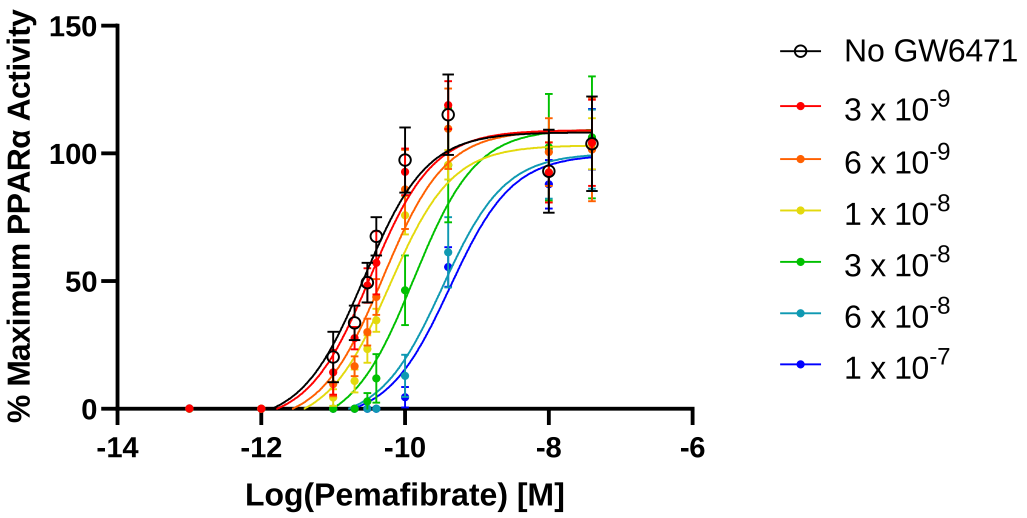

**Figure S2**. Unconstrained pemafibrate dose response curves with increasing concentrations of GW6471. Cos-7 cells were plated in serum-free medium in 96-well plates. Cells were dosed with increasing concentrations of pemafibrate with vehicle (DMSO) or increasing concentrations of pemafibrate with the indicated concentration of the PPARα antagonist GW6471. Luminescence and fluorescence were measured after 24 hrs. Data were calculated as described in the Methods. Data points are the mean and standard deviation of three to four independent experiments. The lowest dose shown for each curve is for wells treated with vehicle only.

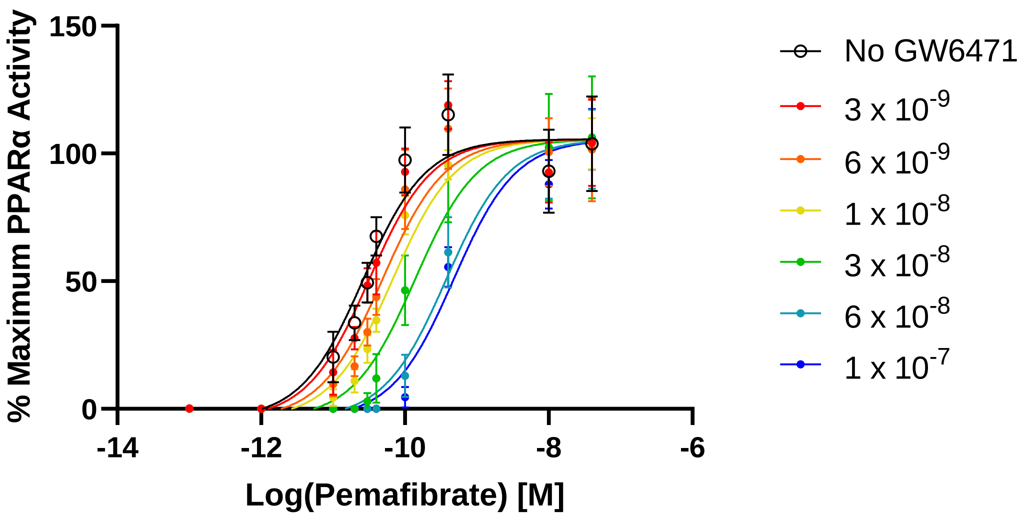

**Figure S3**. Constrained pemafibrate dose response curves with the indicated dose of GW6471. Data generated in cos-7 cells plated in serum free medium. Luminescence and fluorescence were measured after 24 hrs. Data were calculated as described in the Methods. Data for wells treated with vehicle only are shown as the first concentration for each curve. Curve fits were constrained to the mean minimum and maximum of the individual parameters fit in Supplemental Figure 3. Data presented are the mean and standard deviation for three to four replicates.

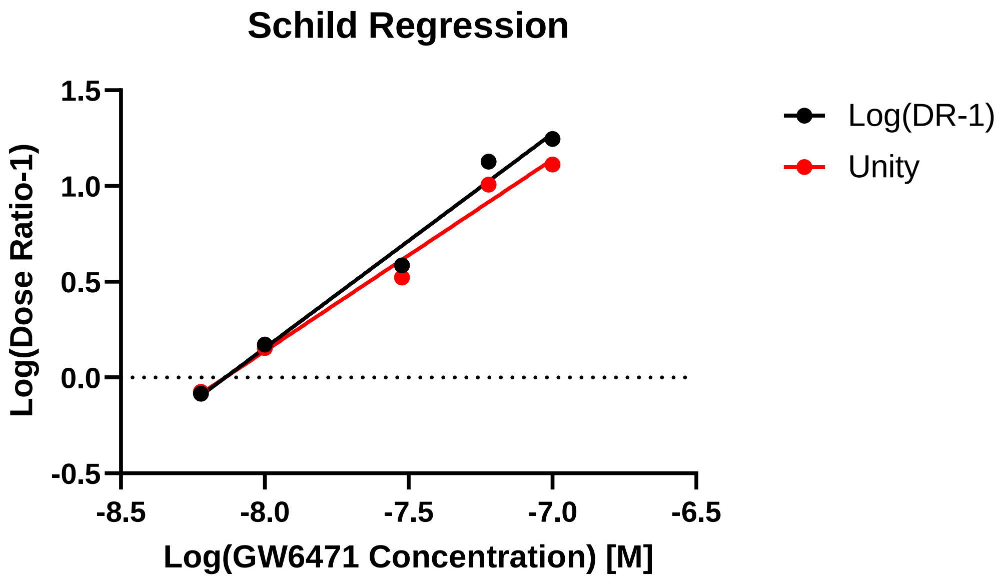

**Figure S4**. Schild regression to estimate equilibrium dissociation constant of GW6471.
